# Rheostatic Network Consolidation Drives Physical Aging in Biomolecular Condensates

**DOI:** 10.64898/2026.06.30.735561

**Authors:** David Polanco, Karinna G. Pele, Alicia Mairo, María Martínez-Monge, Nicolás Moreno, Nunilo Cremades

## Abstract

While the physical aging of biomolecular condensates into macroscopic glasses is heavily linked to pathological disease states, the nanoscale topological rules governing this non-equilibrium relaxation remain elusive. Using heterotypic α-synuclein–Tau coacervates, we combine variable-stringency dissolution and FLIM-FRET to provide direct experimental mapping of the internal network reorganization over time. Rather than a passive, isotropic kinetic jamming event typical of classic glasses, we demonstrate that this physical aging is driven by continuous rheostatic network consolidation; a progressive, directed topological relaxation toward deeper free-energy minima powered by the cooperative spatial optimization of sticker motifs. We formalize these dynamics into a mesoscale series-resistance model derived from size-resolved kinetics, proving that thermodynamic quench depth dictates the initial network state while clustered sticker patterning introduces configurational frustration that kinetically stalls maturation to preserve liquidity. This multi-scale framework links sequence grammar to non-equilibrium transport laws, revealing how biomolecular assemblies navigate the boundary between physiological utility and pathological arrest.

---

Biomolecular condensates organize cellular chemistry across a plethora of biological functions^1–3^ through associative phase separation coupled to percolation, which dynamically generates extensive networks of reversible physical contacts between biomolecules^4–6^. Rather than operating as static equilibrium structures, many condensates are intrinsically metastable, non-equilibrium materials whose internal dynamics progressively slow with time^7–12^. This physical aging does not manifest as an abrupt, binary switch between liquid and solid states, but rather as a progressive transition through a continuum of increasingly dense network-packing states. Recent theoretical paradigms have elegantly captured this macroscopic slowdown by framing biomolecular droplets as aging Maxwell glasses, where constituent elements stochastically navigate a broad landscape of energetic traps, gradually losing their mobility^13,14^. Transitioning from these baseline macroscopic descriptions to a predictive physical framework for non-equilibrium relaxation requires the integration of nanoscale topological reconfigurations of the network with mesoscale transport laws.

The thermodynamics of these architectures have been successfully modeled using the associating polymer theory introduced by Semenov and Rubinstein^15,16^, where interacting heteropolymers are described in terms of “stickers” that associate via specific binding motifs and “spacers” that connect stickers and influence overall solubility, sticker-sticker cooperativity and viscoelastic properties of condensates^9,17,18^. While modern extensions of this theory have elegantly provided a thermodynamic description of associative phase separation coupled to percolation^9,19–21^, these frameworks remain fundamentally tied to equilibrium states. Consequently, temporal material hardening within protein condensates is conventionally ascribed to local conformational changes, particularly the transition of disordered chains into ordered, cross-β amyloid sheets^22–25^. Most mechanistic insights for this derive from homotypic, prion-like low-complexity domains in neurodegeneration-linked proteins, where aromatic and hydrophobic motifs drive both phase separation and amyloid nucleation^9,26–31^. Alternatively, the mean-field models for describing aging in Maxwell glasses^14^ are agnostic to explicit macromolecular architecture, leaving the microscopic drivers of landscape evolution unresolved. Non-equilibrium computational frameworks have proposed a mechanism of valence exhaustion, where kinetic arrest emerges from the progressive, complete saturation of available sticker valencies into long-lived bonds without requiring structural transitions^32,33^. Parallel to this, recent analytical polymer theory suggests that glass-like kinetic trapping can alternatively be driven by the entropic maximization of intervening spacers, which forces stickers into dense, localized clusters over time^34^. Bridging these perspectives, we hypothesize that physical aging in complex heterotypic environments provides the explicit microscopic realization of an aging Maxwell glass through a continuous topological mechanism: rheostatic network consolidation. Under this framework, an initially under-relaxed network progressively deepens its free-energy well by optimizing its spatial connectivity over time.

To test this hypothesis within heterotypic electrostatic networks, whose aging trajectories remain largely unexplored experimentally despite their relevance in a vast subset of cellular functions^35–40^, we established α-synuclein (αSyn) and Tau as a biomedically relevant model system^41,42^. Both form heterotypic fluid coacervates driven by electrostatic complementarity between the anionic C-terminus of αSyn and the cationic motifs of Tau’s central domain (particularly the proline-rich domain, PRD)^43,44^. Crucially, their highly distinct charge architectures—a continuous anionic block in αSyn versus Tau’s segmented, well-dispersed charged motifs—(Fig. 1a,b) provide an ideal orthogonal scaffold to dissect how sequence-level motif patterning and valency govern the ruggedness of the relaxation landscape.

**Fig. 1.**
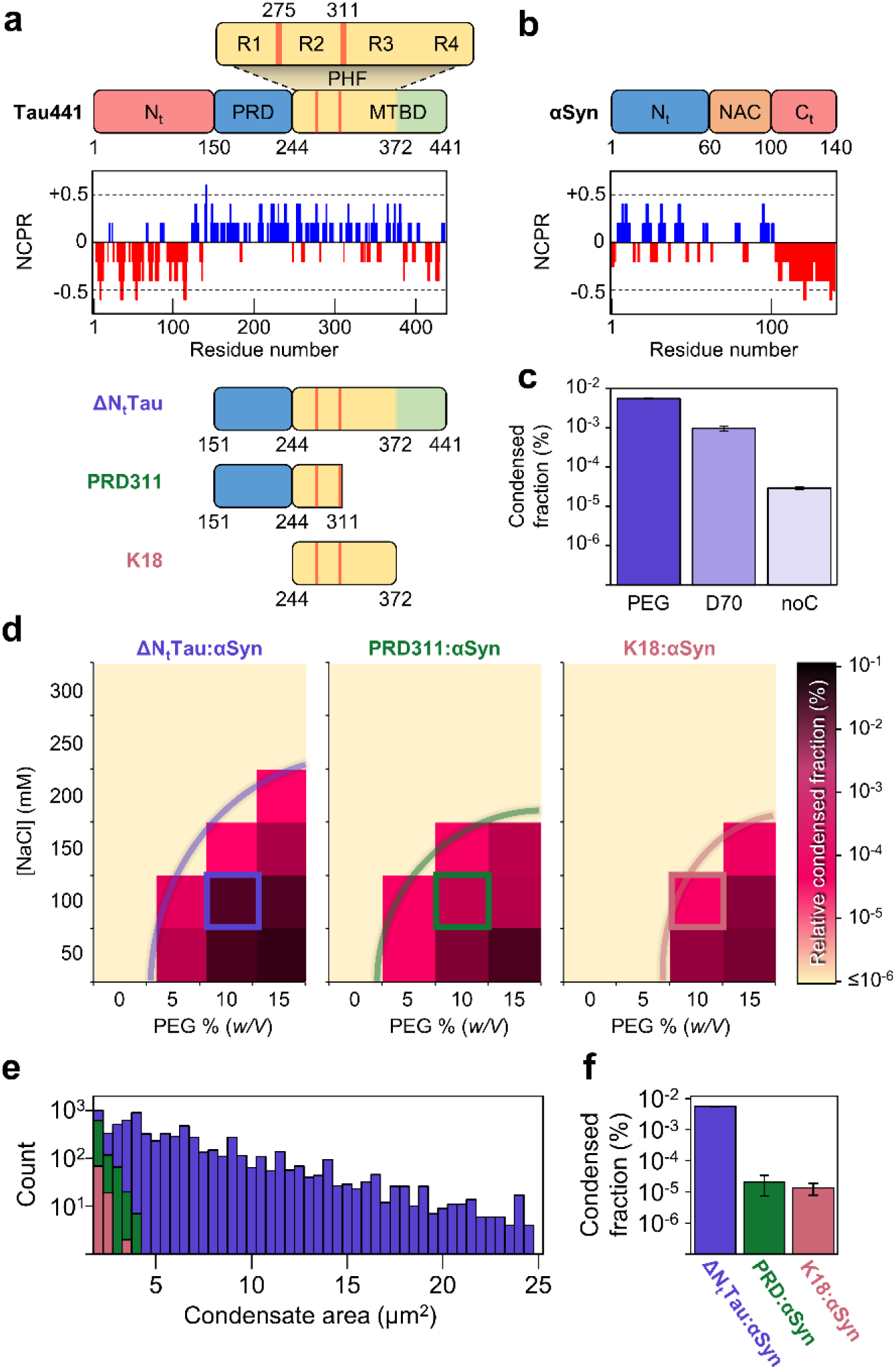
Complex coacervation of αSyn with three Tau constructs. a, Schematic of full-length (2N4R isoform) human Tau and experimental truncates, showing the negatively charged N-terminal domain (N_t_), proline-rich region (PRD) and amyloid-forming paired helical filament (PHF) region within the microtubule-binding domain (MTBD). The net charge per residue (NCPR) is plotted above the Tau441 scheme. **b,** Schematic and NCPR of wild-type human αSyn, highlighting the amphipathic N-terminal domain, the hydrophobic NAC region, and the highly negatively charged C-terminal domain. **c,** Determined condensed fraction of ΔN_t_Tau:αSyn at 50:50 μM (supplemented with 1 μM AF488-αSyn) 1 h after mixing in PS buffer with 10% PEG8k (PEG) or 15% dextran70k (D70), or in absence of crowder in low ionic-strength buffer - 20 mM NaCl (noC). **d,** Phase diagrams mapping the relative condensed fraction of heterotypic ΔN_t_Tau:αSyn, PRD311:αSyn and K18:αSyn condensates (50:50 μM) across a grid of NaCl and PEG8k concentrations (27 fields spanning 10 optical sections per condition were acquired using a 40x objective). A uniform intensity-based thresholding strategy was applied across all micrographs to enable intra-and inter-system comparisons. The color scale denotes the fractional area of the condensed phase relative to the total micrograph area; curves indicate estimated phase boundaries, and the standard experimental PS buffer condition is highlighted. **e,** Representative size distributions of ΔN_t_Tau:αSyn (purple), PRD311:αSyn (green) and K18:αSyn (pink) condensates in PS buffer at 50:50 μM recorded 1 h post-mixing (data aggregated from 9 fields across 10 optical planes). **f,** Quantified condensed fraction for the three stoichiometric systems. Data in **c** and **f** represent mean values derived from three independent biological replicates (3 fields x 10 planes per replicate); error bars denote *s.d*. Scale bars, 20 μm.

Here, we combine variable-stringency dissolution assays with *in situ* single-condensate FLIM–FRET imaging to map the evolving nanoscale topology inside individual heterotypic droplets over time. We demonstrate that these condensates undergo a continuous, amyloid-independent rheostatic network consolidation driven by a time-dependent deepening of their free-energy basin. By formalizing these dynamics into a mesoscale series-resistance model, we show that the physical aging rate is strictly governed by an interplay between the non-equilibrium initial quench depth within the phase diagram and the spatial arrangement of interaction motifs. Ultimately, this work provides a predictive framework linking sequence grammar to non-equilibrium transport laws, offering a blueprint for how biomolecular assemblies navigate the boundary between physiological utility and pathological arrest.

## RESULTS

### Kinetic decoupling between phase separation and network stabilization

To dissect how polymer grammar controls network stabilization, we paired αSyn with three cationic Tau variants lacking the acidic N-terminus (Fig. 1a). ΔN_t_Tau (Tau₁₅₁_–_₄₄₁) serves as a long, high valency scaffold; PRD311 (Tau₁₅₁_–_₃₁₁) isolates the effect of shortened contour length at a comparable valency; and K18 (Tau₂₄₄_–_₃₇₂) isolates the effect of lower effective valency and altered sticker distribution within a short contour length.

Under macromolecular crowding at physiological ionic strength (100 mM NaCl) and a 1:1 molar ratio (50 μM each), phase separation propensity followed the hierarchy ΔN_t_Tau:αSyn > PRD311:αSyn > K18:αSyn (Fig. 1d,f). Under these conditions, we analyzed physical aging using independent, variable-stringency dissolution challenges for the three systems (Fig. 2a,b). While freshly nucleated droplets dissolved completely, all systems progressively developed a graded, condition-dependent resistance to dissolution over time (Fig. 2c–e; see Supp. Fig. 4-5 for independent biological replicates and data reproducibility). Control experiments confirmed that crowding favors initial demixing at physiologically relevant protein and salt concentrations, without driving subsequent time-dependent stabilization (Fig. 1c; see also Supp. Results and Supp. Fig. 1–3).

**Fig. 2.**
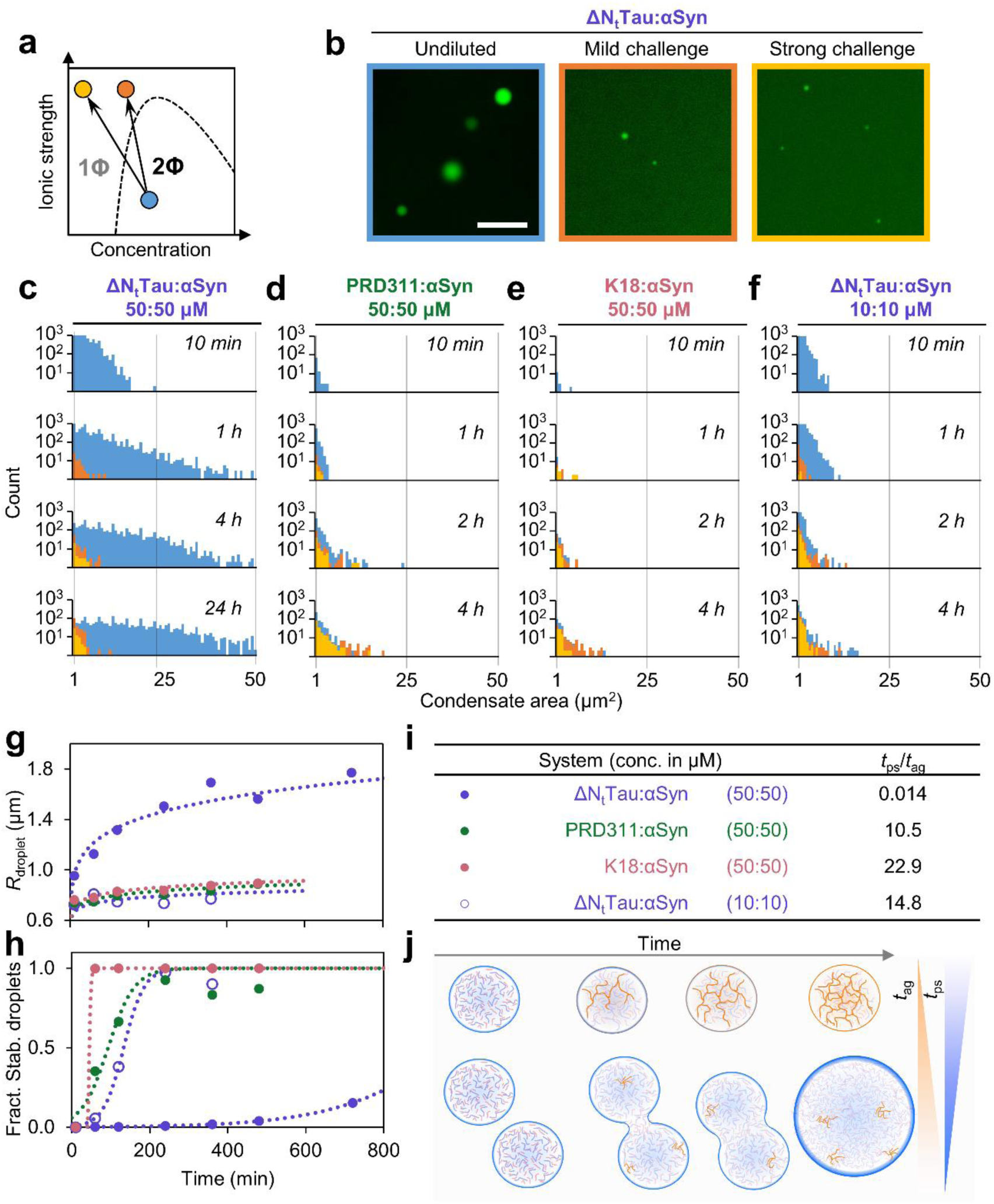
Condensate stabilization kinetics of αSyn with three Tau constructs assessed through electrostatic challenges. a, Schematic phase diagram with a phase boundary (dotted line) separating one-phase (1Φ) and two-phase (2Φ) regimes. Condensate stabilization is evaluated by shifting the initial system state (blue dot) across the phase boundary via low-stringency (orange: elevated ionic strength, mild dilution) or high-stringency (yellow: elevated ionic strength, strong dilution) chemical challenges. **b,** Confocal micrographs of 50:50 μM ΔN_t_Tau:αSyn condensates in PS buffer incubated for 24 h before (undiluted, blue) and after 1:1 dilution into a mild challenge (10% w/V PEG8k, 1 M NaCl; orange) or strong challenge (1 M NaCl; yellow) buffer matrix. Scale bar, 10 μm. **c-f,** Droplet size distributions of ΔN_t_Tau:αSyn (c,f), PRD311:αSyn (d) and K18:αSyn (e) condensates formed in PS buffer and matured at the indicated timepoints. Control distributions in baseline PS buffer are shown in blue; challenged populations are shown in orange (mild) or yellow (strong). Data represent a single representative experimental replicate aggregated from 9 fields across 10 optical sections per longitudinal time point (see Supp. Fig. 3-4 for an inter-replica analysis). **g,h,** Estimated rates of phase separation (g, droplet radius growth) and internal network stabilization (h, stabilized mass fraction against the mild challenge) derived from histogram analysis (*n* = 3): for ΔN_t_Tau:αSyn at 50:50 μM (purple, filled markers), PRD311:αSyn at 50:50 μM (green), K18:αSyn at 50:50 μM (pink) and ΔN_t_Tau:αSyn at 10:10 μM (purple, hollow markers). Dotted curves display numerical fits for radius growth and stabilized-fraction evolution. **i**, Estimated characteristic time constants (*t*_ps_⁄*t*_ag_) against the mild challenge for each independent coacervate system under the experimental conditions tested. **j,** Schematic representing the kinetic competition between condensate stabilization and phase separation characteristic timescales to show how the underlying ratio of macro-phase separation to internal network gelation timescales dictates the morphology and fluid properties of the dense phase, yielding either small gel-like networks or large liquid-like droplets.

Quantifying the apparent kinetic rates and characteristic timescales for phase separation (*k*_ps_, *t*_ps_) and network stabilization/aging (*k*_ag_, *t*_ag_) (see Extended Methods in Supplementary Material) revealed a striking kinetic decoupling (Fig. 2g-i). For ΔN_t_Tau:αSyn system, rapid droplet growth outpaces internal relaxation (*t*_ps_ << *t*_ag_). Conversely, for shorter variants under the same experimental conditions, network stabilization occurs faster than droplet coarsening (*t*_ps_ > *t*_ag_), effectively bounding the ultimate droplet size. Lowering total protein concentration (10:10 μM) reduced the encounter-frequency advantage of the longer ΔN_t_Tau chain, causing its *t*_ps_/*t*_ag_ to converge with the shorter variants. Under these conditions, the apparent rate of stabilization of ΔN_t_Tau:αSyn became remarkably similar to PRD311:αSyn (similar valency), whereas with K18 (lower valency) exhibited the fastest stabilization rate (Fig. 2h). These data indicate that while chain length strongly modulates the macroscale formation and size distribution of condensates, local valency and motif architecture tune the intrinsic rate of internal network maturation. Droplet population and stability therefore emerges from a dynamic competition between condensate growth and internal network stabilization (Fig. 2j).

### Condensate stabilization proceeds independently of amyloid formation

The characteristic timescale of network stabilization (*t*_ag_ ≍ 0.8 h for K18:αSyn, and ≍ 2 h for PRD311:αSyn or ΔN_t_Tau:αSyn) was found to be an order of magnitude shorter than previously reported kinetics for canonical cross-β amyloid assembly in αSyn under amyloid-promoting conditions (*t*_1/2_ ≍ 35 h)^45^. Fluorescence condensate mixing assays further demonstrated that while aging constrained bulk molecular rearrangements (reducing droplet mixing from 88 ± 6 % in freshly prepared samples to 45 ± 9 % in consolidated K18:αSyn condensates; Fig 3g and Supp. Fig. 6), it preserved residual internal mobility rather than inducing irreversible, rigid aggregation. Furthermore, aged, dissolution-resistant condensates exhibited no detectable fluorescence signal from the amyloid-sensitive dye Amytracker-630 (Fig. 3a-c), dissolved completely under conditions known to leave amyloid structures intact (2% SDS, 0.5 M GdnHCl)^46,47^ (Fig. 3f), and retained identical aging kinetics when evaluated using aggregation-deficient variants lacking core amyloidogenic motifs (Δ(72-81)αSyn – referred to as AggDef-αSyn – and Tau-PRD273) (Fig. 3d,e). Consequently, condensate maturation reflects a progressive reorganization of the non-equilibrium electrostatic network rather than cross-β sheet nucleation.

**Fig. 3.**
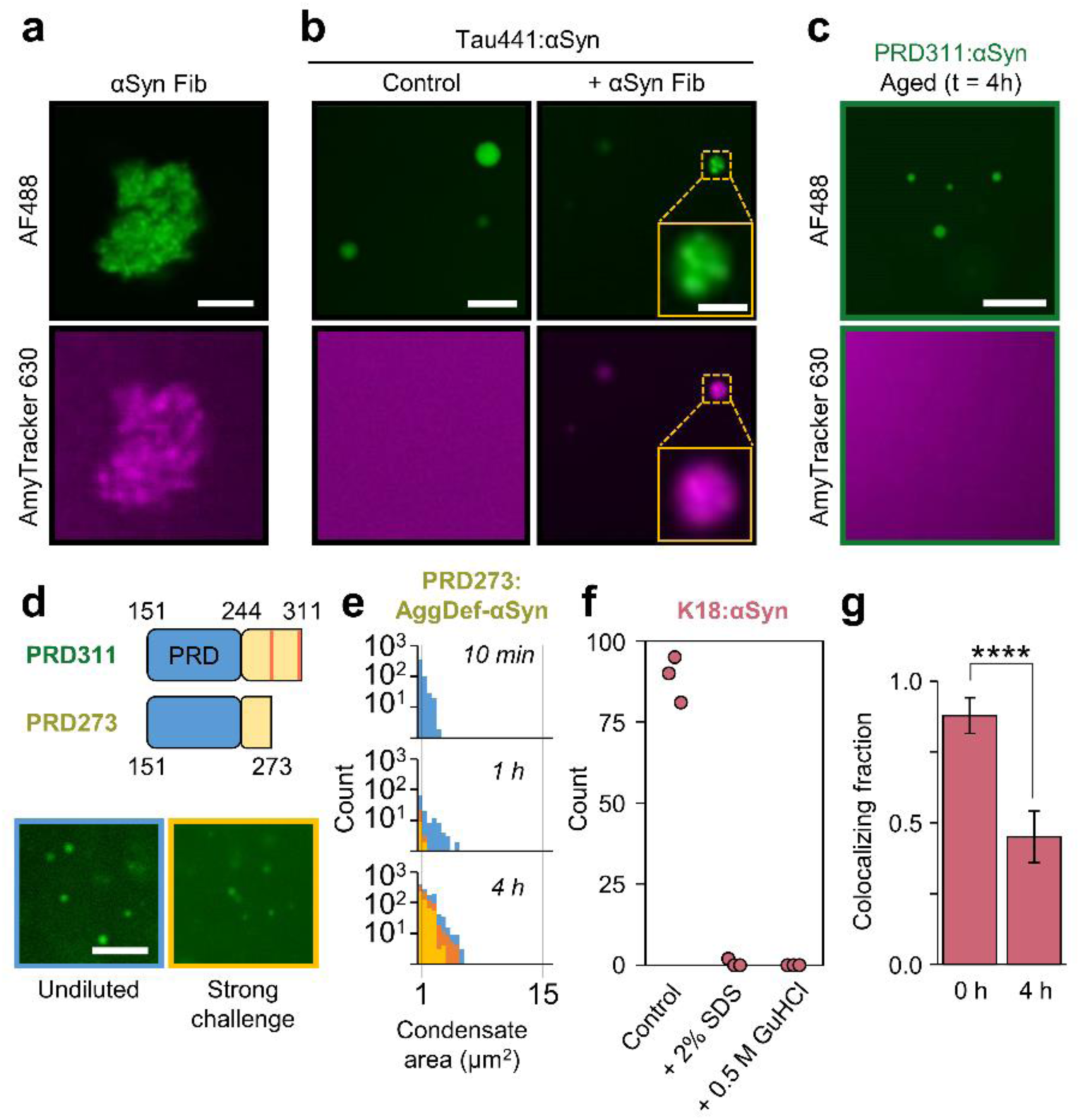
Condensate stabilization differs from amyloid aggregate formation. a, Representative confocal image of unsonicated αSyn fibrils (20% AF488-αSyn labeled, green) 10 min post-addition of 1 μg/mL Amytracker 630 (magenta). **b,** Confocal images of 10:10 μM Tau441:αSyn condensates (supplemented with 1 μM AF488-αSyn, green) 1 h post-formation in PS buffer, without (Control) or with sonicated pre-formed αSyn amyloid fibrils (20% AF488-labelled αSyn fibrils). Samples were imaged across both fluorophore channels 10 min post-addition of 1 μg/mL Amytracker 630 (magenta), revealing that exogenous αSyn fibrils preferentially partition into the fluid coacervate phase. The yellow inset highlights a zoomed condensate with embedded αSyn fibrils. **c,** Confocal images of 50:50 μM PRD311:αSyn (supplemented with 1 μM AF488-αSyn, green) condensates 4 h post-formation in PS buffer and 10 min post-addition of Amytracker 630 (magenta). **d,** Structural schematic of PRD311 and the aggregation-deficient PRD273, which lacks the amyloidogenic hexapeptide motifs (red lines). Confocal images show 50:50 μM PRD273:αSyn condensates (supplemented with 1 μM AF488-αSyn) after 4 h incubation before (undiluted, blue) and after low-stringency (mild, orange) or high-stringency (strong, yellow) chemical challenges. Scale bars: 10 μm for main panels a-d, 2 μm for insets in b. **e,** Longitudinal droplet size distributions for 50:50 μM PRD273:αSyn condensates (baseline, blue; mild challenge, orange; strong challenge, yellow). **f,** Total remaining droplet count of 50:50 μM K18:αSyn condensates after 24 h incubation followed by 1:1 high-stringency challenge and subsequent treatment with 2% (w/V) SDS or 0.5 M guanidinium hydrochloride. **g,** Colocalizing fraction measured 1 h post-mixing of separately prepared K18:αSyn condensates (50:50 μM; labeled with either 1 μM AF488-αSyn or Atto647N-αSyn) in PS buffer. Mixing pre-formed condensates of each color was performed either immediately after phase separation (0 h) or following a 4 h pre-incubation period. Bars denote the mean colocalizing fraction computed from 6 independent fields across 10 optical sections (*n* = 526 droplets for 0 h; *n* = 192 droplets for 4 h); statistical significance was determined by a two-tailed Student’s *t*-test (p = 2.85 x 10^-6^).

### Single-condensate FLIM-FRET resolves rheostatic network consolidation *in situ*

To track nanoscale restructuring, we developed an *in situ* single-condensate FLIM-FRET assay utilizing AF488 as donor on the acidic C-terminus of αSyn (position 122) and a non-fluorescent quencher (TQ2WS) on the cationic domains of Tau. Progressive decreases in donor lifetime in the presence of the quencher (+Q), compare to its absence (-Q), directly reported a time-dependent increase in proximity and cooperative strengthening between interacting motifs, tracking macroscopic dissolution resistance (Fig. 4a,b).

**Fig. 4.**
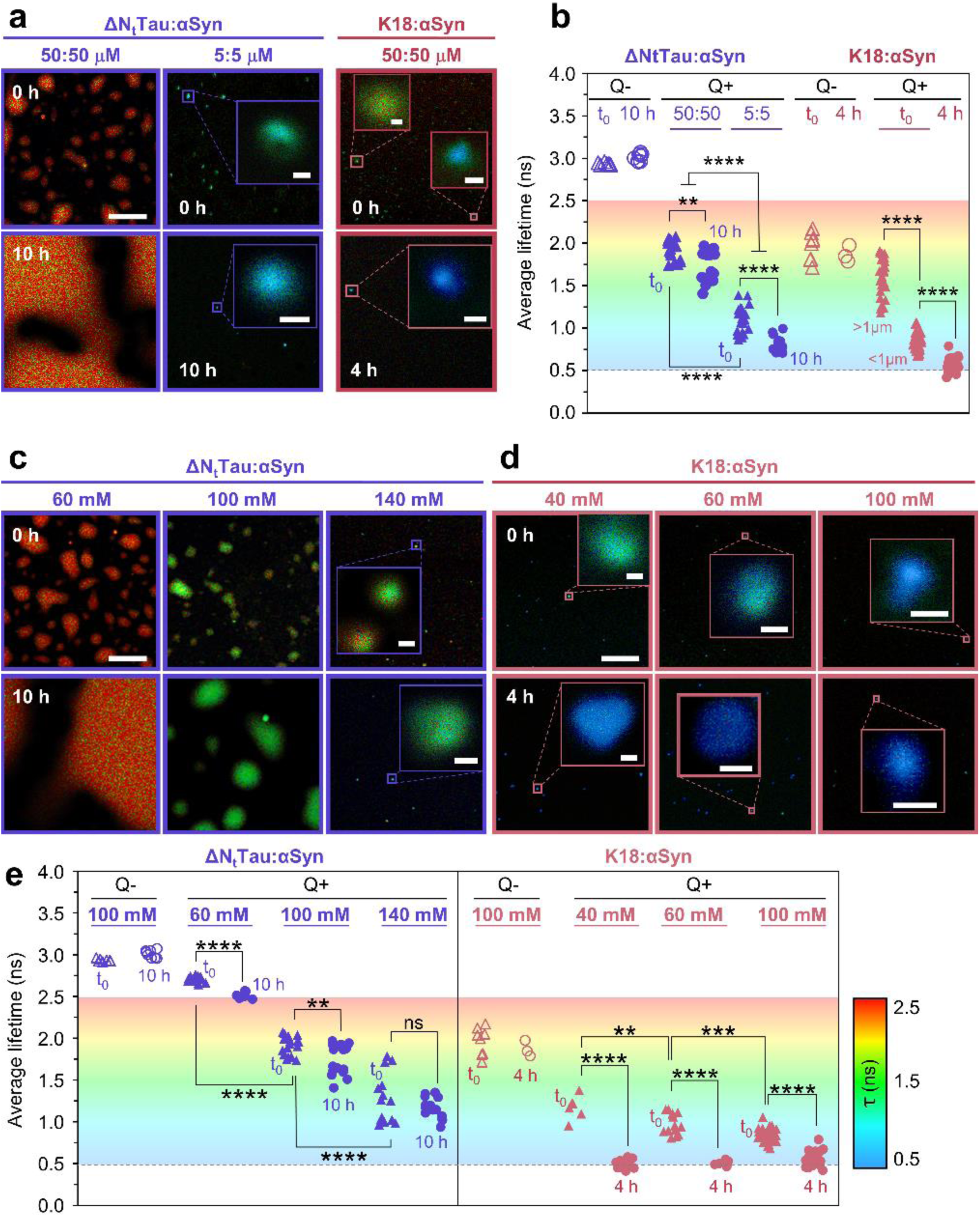
Condensate stabilization kinetics depends on quench depth of the initial network. a, Representative FLIM images showing the intensity-weighted average AF488 lifetime in FRET-active ΔN_t_Tau:αSyn condensates (purple; formed at 50:50 μM [left], or 5:5 μM [right] in PS buffer) and K18:αSyn condensates (pink; formed at 50:50 μM in PS buffer). FRET pairs comprise a substoichiometrically C-terminally labeled AF488-αSyn donor (5-10% of total αSyn concentration) and a Tau acceptor bearing a TQ2WS quencher on its positively charged stickers (50% of total Tau concentration). Lifetimes were recorded immediately post-formation (0 h) and following a 10 h (ΔN_t_Tau:αSyn) or 4h (K18:αSyn) incubation. **b,** Absolute AF488 lifetimes in FRET active (Q+) ΔN_t_Tau:αSyn condensates (purple) formed at 50:50 or 5:5 μM in PS buffer measured at 0 h (triangles) or 10 h (circles), and in Q+ K18:αSyn condensates (pink) formed at 50:50 μM measured at 0 h (triangles) or 4 h (circles). Donor-only control (Q-) lifetimes lacking the TQ2WS acceptor quencher at 50:50 μM are shown as hollow markers. **c,** Representative FLIM images of FRET active ΔN_t_Tau:αSyn condensates formed at 50:50 μM in PS buffer containing 60, 100 or 140 mM NaCl, measured at 0 h and 10 h. **d,** Representative FLIM images of FRET active K18:αSyn condensates formed at 50:50 μM in PS buffer containing 40, 60 or 100 mM NaCl, measured at 0 h and 4 h. **e,** Absolute AF488 lifetimes in Q+ ΔN_t_Tau:αSyn (purple) and K18:αSyn (pink) condensates across the designated ionic strengths, measured at 0 h (triangles) and 10 h or 4 h (circles). Lifetimes for Q-control condensates at 100 mM NaCl are shown as hollow markers. a theoretical minimum lifetime for the compacted K18:αSyn network is indicated by a horizontal dotted line at 0.5 ns. Scale bars: 20 μm for main panels; 500 nm for insets. Color scales indicate average lifetime per pixel; background gradients match the pseudocolor lifetime scale to indicate continuous transitions across samples for all panels.

K18:αSyn exhibited tighter initial molecular packing and accelerated compaction compared to ΔN_t_Tau:αSyn. Intradroplet analysis of K18:αSyn revealed a pronounced size dependence: smaller droplets consistently showed shorter donor lifetimes and faster relaxation rates than larger ones. Similarly, ΔN_t_Tau:αSyn condensates formed at low protein concentrations exhibited shorter initial lifetimes, which decreased more sharply after 10 h of maturation than condensates at higher protein concentrations (Fig. 4b). This network compaction was strictly regiospecific; donor labeling at the distal N-terminus of αSyn (position 24) yielded negligible lifetime changes over time in ΔN_t_Tau:αSyn and K18:αSyn droplets (Supp. Fig. 7; see also Supp. Results), proving that aging is driven by the localized spatial proximity of specific electrostatic motifs rather than a non-specific, global collapse of the polypeptide chains.

### Quench depth sets the initial network state and heterogeneity

Modulating solution ionic strength decoupled the initial network topology from polymer identity. Increasing NaCl shifted the system toward the binodal boundary (shallow quench), producing smaller droplets born with short donor lifetimes already close to their relaxed configuration (Fig. 4c–e), mirroring the effects of lowering protein concentration. Conversely, decreasing NaCl drove a deep quench into the two-phase regime, triggering explosive droplet growth that initially captured chains within large, under-relaxed network topologies characterized by long fluorescence lifetimes, which subsequently shortened over time during structural relaxation. These findings indicate that the non-equilibrium depth of the initial thermodynamic quench dictates the starting configurational state of the network.

Remarkably, droplet-to-droplet FLIM-FRET heterogeneity peaked sharply during intermediate network consolidation regimes before ultimately converging toward a uniform, limiting lifetime configuration, regardless of the initial network state of the nascent condensates (Fig. 4e). This transient structural divergence followed by strict convergence provides direct coordinate-level evidence that physical aging represents a directed, progressive consolidation of the interaction network toward a structurally optimized free-energy minimum. This multi-scale maturation is coordinated by localized network reconfigurations that macroscopically manifest as condensate shrinking^8^, the cooperative formation of transient nanoscale high-density domains^48^, and the concomitant expulsion of solvent^49,50^. Crucially, these data demonstrate that biomolecular condensates do not simply toggle between binary, discrete liquid and gel phases; rather, they populate a continuous, rheostatic spectrum of network-packing states as they navigate a rugged but decisively funneled relaxation landscape.

### Sticker architecture sets the kinetic bottleneck for aging

To determine how sequence grammar guides landscape navigation, we contrasted the Tau variants against the C-terminal domain of histone H1 (CH1), an IDR of similar size to PRD311 but featuring a high, uniform charge density arranged into a nearly continuous sticker motif. Remarkably, CH1:αSyn condensates completely failed to acquire dissolution resistance after 24 hours under identical conditions to the aging Tau:αSyn networks (Fig. 5a and Supp. Fig. 8). This demonstrates that unpatterned charge distributions cannot sustain productive network optimization; instead, it remains trapped in an under-relaxed, highly reversible state, aligning with polymer mean-field models where high charge disparity without discrete topological organization precludes efficient network percolation^51^.

**Fig. 5.**
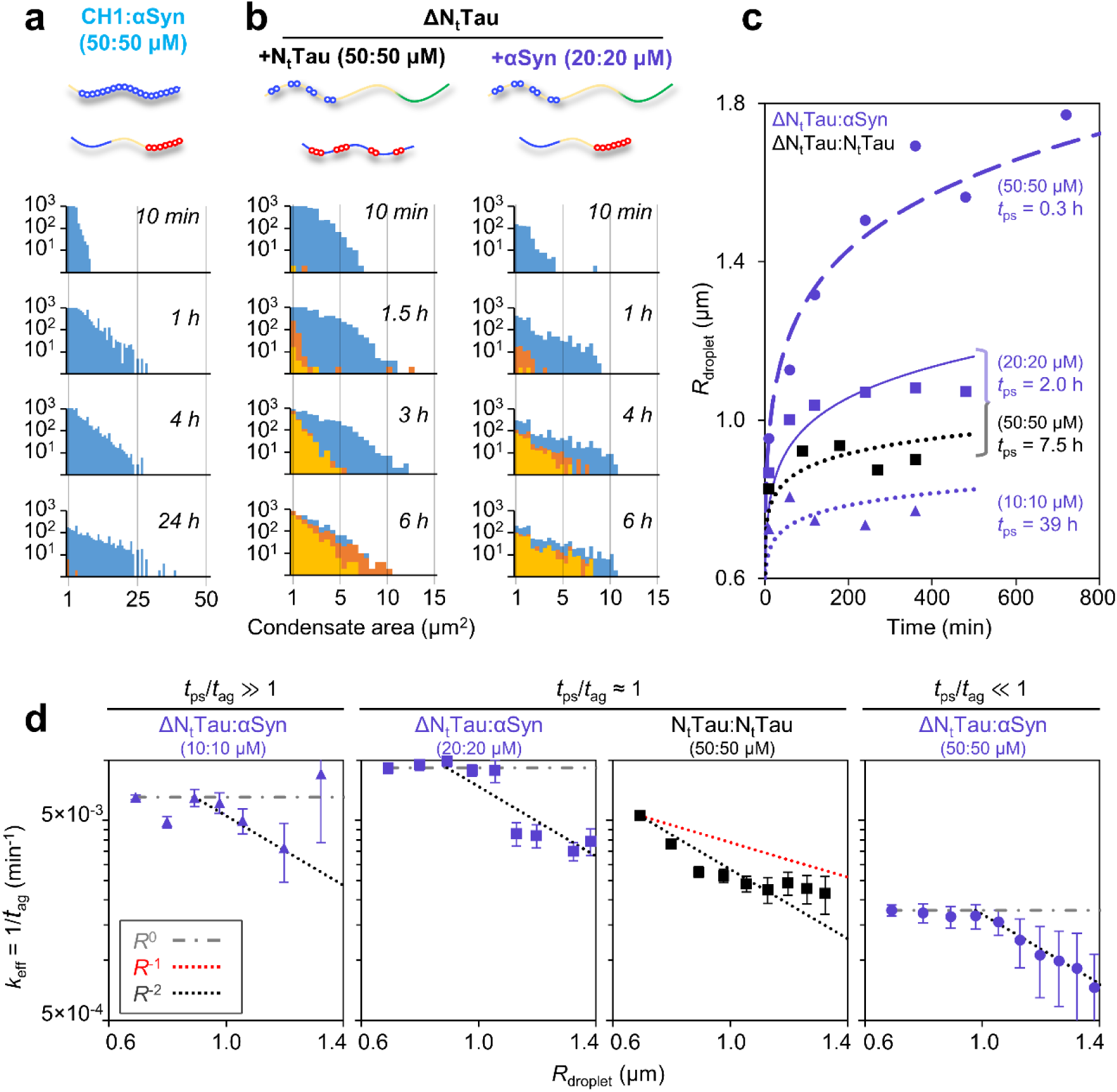
Condensate stabilization kinetics depend on polymer architecture: well-dispersed vs continuous stickers. a, Longitudinal droplet size distributions for 50:50 μM CH1:αSyn condensates (supplemented with 1 μM AF488-αSyn) in PS buffer and imaged at the indicated time points (baseline, blue; mild challenge, orange; strong challenge, yellow). Data represent a single representative replicate aggregated from 9 fields across 10 optical slices. **b,** Size distributions of 50:50 μM ΔN_t_Tau:N_t_Tau (left) and 20:20 μM ΔN_t_Tau:αSyn (right) condensates (each containing 1 μM of the respective fluorescently labeled, negatively charged partner) in PS buffer. Representative replicates comprise 18 fields across 16 optical sections per time point. **c,** Rates of macro-phase separation (radius growth) for ΔN_t_Tau:N_t_Tau at 50:50 μM (filled black squares, solid fit) and ΔN_t_Tau:αSyn (purple) at 50:50 μM (filled circles, dashed fit), 20:20 μM (filled triangles, solid fit), and 10:10 μM (hollow circles, dotted fit). The similar phase-separation characteristic timescales shared by 50:50 μM ΔN_t_Tau:N_t_Tau and 20:20 μM ΔN_t_Tau:αSyn condensates allow a direct comparison between effective network consolidation rates. **d,** Estimated size-resolved effective stabilization *k*_eff_ rate constants (referred to as *N*_drop_) as a function of droplet radius against low-stringency (mild) challenge for the systems in **c** (see Supp. Fig. 8 for high-stringency challenge analysis). Points denote experimental values derived from the mean of three independent biological replicates; error bars denote relative uncertainty in *k*_eff_ for each droplet size cohort, estimated via Poisson-like counting statistics as σ_*k*eff_⁄*k*_eff_= 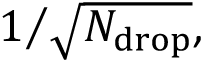 reflecting the finite number of droplets (*N*_drop_) sampled per radius bin; dotted lines serve as visual guides for *R*^0^, *R*^−1^ and *R*^−2^ scaling.

Conversely, pairing ΔN_t_Tau with its endogenous partner N_t_Tau, which features well-dispersed, discrete negatively charged stickers separated by spacers, led to rapid stabilization (Fig. 5b, left panel). To decouple phase separation from network consolidation, we compared ΔN_t_Tau:N_t_Tau (50:50 μM) and ΔN_t_Tau:αSyn (20:20 μM) under concentrations regimes tailored to yield similar initial phase-separation timescales (*t*_ps_) (Fig. 5b,c). Analyzing the network stabilization rate as a function of droplet radius (*R*) (defined here as the size-resolved effective rate, *k*_eff_ = 1/*t*_ag_) revealed distinct scaling regimes (*k*_eff_ ∝ *R*^α^): for ΔN_t_Tau:N_t_Tau, *k*_eff_ scaled inversely with droplet radius across all analyzed sizes, whereas for ΔN_t_Tau:αSyn, *k*_eff_ remained size-independent in small droplets before decaying past a clear crossover radius (Fig. 5d and Supp. Fig. 8).

We formalized these dynamics using a minimal mesoscale series-resistance model, where the net consolidation pathway proceeds through three sequential steps (see theoretical framework descriptions in Supp. Material): 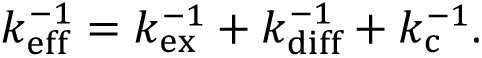 Here, *k* _ex_ represents the interfacial exchange rate of proteins between the phases (scaling with *R*^α^, *α* = −1), *k*_diff_ is the intra-condensate diffusive mixing rate (α = −2), and *k*_c_ is the size-independent molecular consolidation step (α = 0), reflecting the intrinsic rate at which transient intermolecular contacts reorganize into stable cross-links. The slowest step dictates the dominant regime.

This framework reveals that sticker architecture fundamentally alters network topology and the underlying network consolidation free-energy barrier. In the well-dispersed ΔN_t_Tau:N_t_Tau system, a high encounter probability facilitates rapid percolation; *k*_c_ is fast, meaning aging is exclusively transport-limited and size-dependent. In contrast, the clustered, near-continuous anionic architecture of αSyn C-terminal region introduces sliding constraints, steric hindrance and topological frustration. This configurational frustration drastically depresses *k*_c_, rendering it the primary, size-independent rate-limiting step (α ≈ 0) in small droplets. As droplets grow, transport resistance eventually overtakes consolidation, generating the crossover size beyond which the aging rate decays (α < 0) (Fig. 5d). Small condensates thus report on the chemistry of network formation (*k*_c_) for frustrated motifs or transport rates (*k*_ex_ and/or *k*_diff_) for polymers with dispersed stickers.

Physically, *k*_c_ encodes the kinetic barrier required for local contact unbinding and re-alignment proportional to the flow activation energy^52^. Intriguingly, the apparent *k*_c_ for ΔN_t_Tau:αSyn was almost an order of magnitude slower under deep quenches (50:50 μM) than under shallow quenches (20:20 μM or 10:10 μM) (Fig. 5d; Supp. Fig. 9). Because shallow quenches nucleate nascent networks closer to their global thermodynamic minimum, they experience a significantly lower activation barrier to reorganize, as also observed in FLIM-FRET assays (Fig. 4a, ΔN_t_Tau:αSyn system). The non-equilibrium quench depth thus fundamentally modulates the activation energy barrier, governing subsequent network consolidation.

## DISCUSSION

This study identifies a molecular and mesoscale mechanism for the physical aging of biomolecular condensates, using heterotypic αSyn:Tau coacervates to uncover universal principles of network consolidation. The central conclusion is that aging is a continuous, time-dependent relaxation of an initially under-relaxed network, progressing via rheostatic network consolidation rather than an abrupt, binary liquid-solid switch. While macroscale models frame this continuous trajectory as an aging Maxwell glass navigating sequence-agnostic energetic traps^14^, our work provides the first direct experimental visualization of the evolving nanoscale structural topology driving this continuum, demonstrating that macroscopic consolidation can proceed via a purely topological, amyloid-independent pathway. By leveraging *in situ* single-condensate FLIM–FRET, we map the progressive optimization of heterotypic molecular contacts, demonstrating three major conceptual advances.

First, our *in situ* FLIM–FRET and variable-stringency dissolution data provide a quantitative bridge between macroscale glass-like aging and microscopic network consolidation. We show that heterotypic condensates are born into non-equilibrium network topologies strictly dictated by the initial quench depth within the phase diagram, subsequently consolidating over time as sticker motifs achieve greater spatial proximity and cooperatively strengthen. Mechanistically, this progression is consistent with either spacer-driven entropic clustering^34^ or a locally propagated valence exhaustion cascade^10^. Crucially, our sequence-patterning variations demonstrate that regardless of the underlying microscopic driver, a clustered sticker architecture introduces configurational frustration that stalls network consolidation.

Second, these results extend recent models characterizing the thermodynamic relationship between condensate and amyloid formation. Beyond serving as interfaces that promote nucleation^29,30,53,54^ or interiors that buffer aggregation^30^, aging condensates undergo progressive energy basin deepening into an optimized thermodynamic sink, continuously lowering their internal chemical potential. As the network tightens, its free-energy well deepens, raising the activation energy barrier for structural reorganization into ordered, crystalline β-sheets^44^. Physical aging thus acts as a temporary kinetic shield, trapping the assembly in a stable amorphous state and delaying conversion to the global amyloid minimum. Consequently, amyloid risk depends fundamentally on where an assembly resides along its internal relaxation landscape (Fig. 6a,b).

**Fig. 6.**
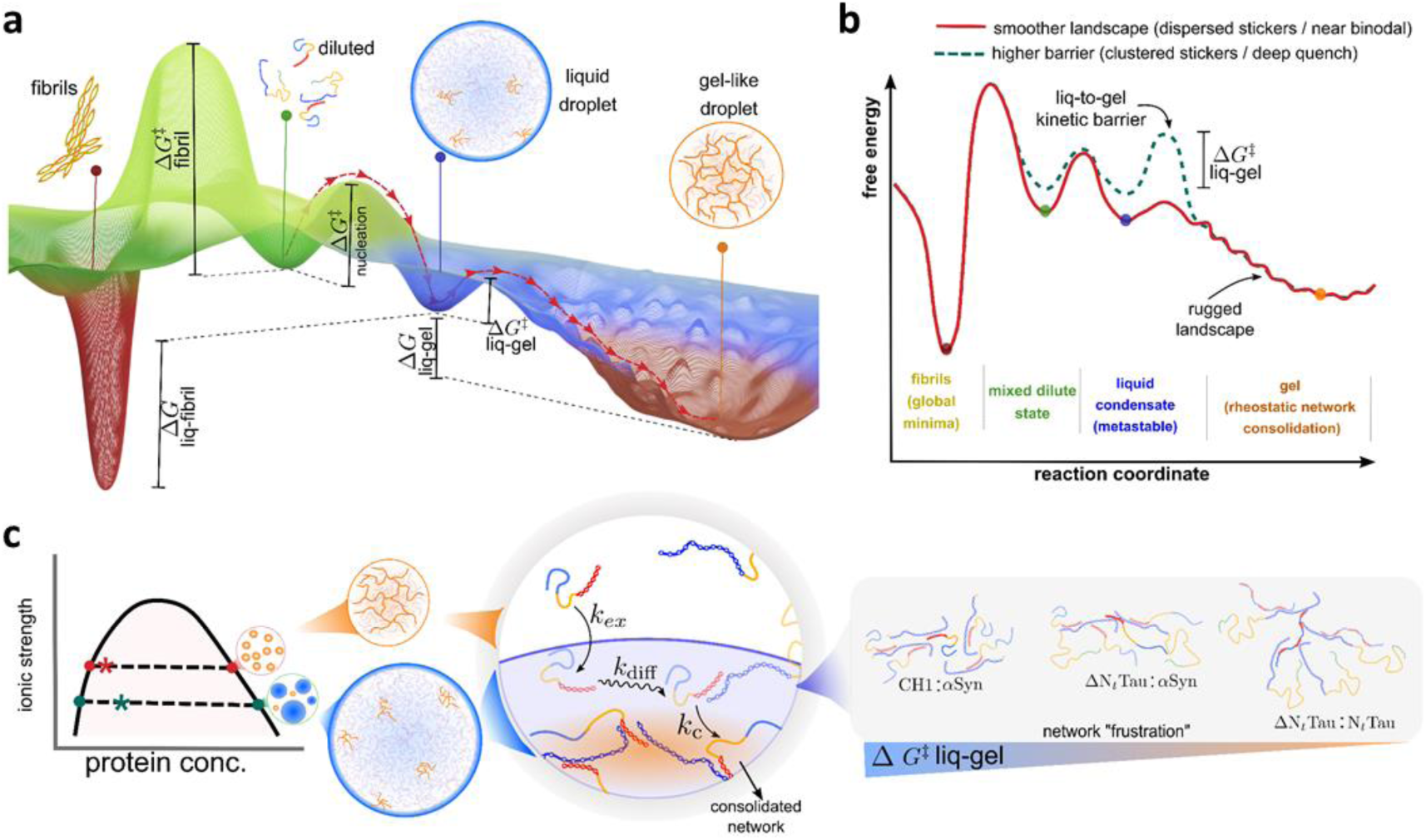
Free-energy landscapes and kinetic routes governing heterotypic coacervation, gelation, and fibrillation. a, Schematic multidimensional free-energy landscape for the assembly of a two-protein heterotypic system. Soluble, mixed protein molecules occupy a shallow basin and undergo phase separation by crossing a nucleation barrier (ΔG^‡^ nucleation) to form a liquid-like condensate. From this metastable intermediate, the system can evolve into an amorphous gel via dense-phase network consolidation over a liquid-to-gel barrier (ΔG^‡^ liq-gel). Alternatively, protein molecules in the diluted phase can nucleate into an ordered amyloid state via a separate fibrillation barrier (ΔG^‡^ fibril). The depths of minima and barrier heights determine whether condensates remain liquid, consolidate as gel-like condensates or produce fibrillar assemblies. **b,** One-dimensional landscape projection showing how interaction topology and quench depth reshape aging pathways. Near the binodal, well-dispersed sticker interactions generate a smooth landscape (solid red curve); rather than a discrete, barrier-limited transition, condensates relax through a continuum of increasingly consolidated network states via rheostatic network consolidation within a rugged free-energy funnel. Deep quenches or continuous sticker clustering reshape this into a highly rugged landscape (dashed green curve), elevating ΔG^‡^ liq-gel and trapping metastable liquid droplets. **c,** Molecular determinants governing resistance to network consolidation. The liquid-to-gel transition reflects an underlying kinetic competition among material exchange with the dilute phase (*k*_ex_), internal diffusion-driven rearrangements (*k*_diff_) and network crosslinking (*k*_c_). Proximity to the non-equilibrium phase boundary preserves high molecular mobility, promoting network reorganization and reducing kinetic frustration. Conversely, strong thermodynamic driving forces or clustered sticker architectures accelerate kinetic frustration. This increased configurational frustration raises ΔG^‡^ liq-gel barrier, slows structural equilibration and modulates whether the dense phase remains fluid, becomes viscoelastic, or progresses to solid-like fibrillar assemblies.

Third, we demonstrate that the spatial arrangement of amino acid motifs encodes the precise kinetic bottleneck governing the aging rate by modulating the ruggedness of the network consolidation landscape (Fig. 6c). While sticker identity and valency control phase boundaries^5,9,55^, patterning dictates how an under-relaxed network efficiently navigates its non-equilibrium landscape (Fig. 6b). Well-dispersed complementary motifs create a smooth energy funnel that minimizes the activation energy barrier for rapid network consolidation. In contrast, clustered architectures introduce profound configurational frustration, manifesting as an increased activation barrier for cooperative reorganization. This frustration traps the under-relaxed network in local metastable traps^14^, markedly reducing the molecular consolidation rate (*k_c_*) and explaining why intrinsically disordered regions (IDRs) with high nominal valency can promote phase separation yet stall material maturation. Persistent liquidity, thus, emerges from sequence-encoded topological frustration that prevents the network from escaping local traps to find its consolidated ground state.

By formalizing these dynamics into a mesoscale series-resistance model, we establish a predictive framework that successfully decouples size-independent chemical consolidation from size-dependent transport limitations (Fig. 6c). By treating the consolidating network as a variable resistor whose internal transport resistance scales dynamically, our model demonstrate that physical aging appears size-independent when molecular consolidation (*k*_c_) acts as the slow, rate-limiting bottleneck, but transitions to a size-dependent regime when macroscopic transport-limited steps dominate network consolidation. Crucially, the non-equilibrium quench depth does not merely dictate the initial thermodynamic state of the dense phase (Fig. 6c); it fundamentally modulates the activation free energy barrier for subsequent network stabilization (Fig. 6b). Ultimately, mapping the interplay between sequence-level motif architecture and thermodynamic quench depth enables the accurate prediction of a condensate’s material lifetime by isolating the dominant kinetic resistance along its consolidation pathway.

These principles bear profound implications for the crowded cellular environment. Our finding that near-boundary, shallow quenches yield more compact nascent networks that consolidates rapidly suggests that subtle homeostatic shifts can trigger non-linear changes in condensate lifetimes. Cells may actively exploit sequence-encoded configurational frustration as a protective mechanism to preserve the dynamic, liquid-like utility of multivalent assemblies and prevent premature gelation. In neurodegenerative contexts, mutations or aberrant post-translational modifications affecting αSyn and Tau charge distribution, stoichiometry, or quench depth may smooth the relaxation landscape, shifting condensates into long-lived, hyper-consolidated states.

Several critical limitations outline immediate next steps. While our model aligns with the regulated, time-dependent stabilization recently observed in cytoplasmic TDP-43 assemblies^12^, it must be validated within compositionally complex environments. Additionally, although our *in situ* FLIM–FRET successfully resolved nanoscale structural compaction, pairing this approach with active oscillatory microrheology or optical tweezers will be invaluable to map these structural packing states to explicit changes in viscoelastic moduli. Finally, quantifying the exact thermodynamic energy gap separating the fully consolidated condensate state from the global amyloid minimum, as well as determining how this compacted topology alters long-term interfacial nucleation, remains essential to bridge early-stage network consolidation landscape with emerging “sink-interface” models of biomolecular assemblies^30,56–59^. In this context, mesoscale computational approaches capable of simultaneously capturing aging, coarsening, and phase-separation dynamics across extended spatiotemporal scales^60,61^ will be instrumental in linking molecular-scale structural transitions to emergent condensate behavior.

In summary, we have demonstrated that the physical aging of biomolecular condensates is governed by an interplay between sequence-level molecular grammar and mesoscale transport. Condensates nucleate as under-relaxed networks that behave as aging Maxwell glasses. Their subsequent lifetime is determined by a hierarchy of kinetic resistances, dictated by a quench-depth-modulated activation barrier, droplet size, and the structural capacity of their constituent stickers to overcome configurational frustration. By bridging the nanoscale proximity and cooperative strengthening of interaction motifs with macroscopic material stability via rheostatic network consolidation, this work provides a robust, predictive, and universal framework for understanding how and when biomolecular condensates transition from dynamic functional assemblies into arrested, pathologically vulnerable states.

## METHODS

### Protein expression, purification and labeling

Full-length (2N4R isoform) human Tau (Tau441, expression plasmid obtained from Addgene #16316) and all its variants (generated by IVA cloning, primers are specified in Supplementary information, Table 1) were expressed in BL21 *Escherichia coli* strain and purified as described in KrishnaKumar et al.^62^ with the modifications outlined in Gracia & Polanco et al.^44^. The storage buffer for all Tau variants was 20 mM HEPES, 2 mM DTT, 1 mM EDTA, 0.2 mM MgCl_2_, 1 mM PMSF, 50 μM benzamidine, 100 μM leupeptin and 100 mM NaCl, except for Tau441, which was kept at a higher ionic strength of 500 mM NaCl to avoid spontaneous phase separation during freezing/thawing steps.

The N-terminal fragment of Tau (N_t_-Tau) was treated differently due to its negative net charge. After the lysate was cleared by ultracentrifugation, nucleic acids were precipitated with streptomycin sulfate (10 mg/mL) followed by centrifugation. The clarified supernatant was dialyzed against dialysis buffer (20 mM HEPES pH 6.8, 50 mM NaCl, 1 mM EDTA, 0.1 mM PMSF, 50 μM benzamidine, 100 μM leupeptin, 1 U/mL benzonase) and loaded onto an anion-exchange column (HiTrap SPFF, Cytiva, MA, USA) equilibrated in the same buffer. N_t_-Tau was eluted using stepwise increases in NaCl concentration up to 20% elution buffer (dialysis buffer supplemented with 1 M NaCl). Fractions containing N_t_-Tau were identified by SDS-PAGE, pooled, concentrated, and further purified by size-exclusion chromatography on a Superdex 75 pg column equilibrated in storage buffer (10 mM HEPES pH 7.4, 100 mM NaCl, 1 mM PMSF, 50 μM benzamidine, 100 μM leupeptin). Peak fractions were pooled, flash-frozen in liquid nitrogen, and stored at −80 °C. Protein concentration was determined spectrophotometrically.

Wild-type human alpha-synuclein (αSyn, pT7-7asyn WT expression plasmid), the N122C and Q24C variants, and the aggregation deficient variant AggDef-αSyn (Δ(72-81)-αSyn); generated by IVA cloning, primers specified in Table 1) were expressed in BL21 *E. coli* and purified as described in Hoyer et al.^63^,with the modifications specified in Camino et al.^64^. 5 mM DTT were included in all purification steps for the N122C and Q24C αSyn variants to prevent intermolecular disulfide bridge formation.

The C-terminal fragment (residues 112-225) of chicken Histone 1.11 (CH1) in a pET13a plasmid (kindly provided by Prof. Katherine Stott, University of Cambridge) was transformed into E. coli BL21(DE3) competent cells. Cultures were grown in LB medium at 37 °C to an OD_600_ of 0.6–0.7, and expression was induced with 0.5 mM IPTG for 3 h at 37 °C. Purification protocol was based on Gerchman et al.^65^. Briefly, cell pellets were resuspended in lysis buffer (20 mM HEPES pH 6.8, 1 M NaCl, 1 mM EDTA, 0.5 mM PMSF, 50 μM benzamidine, 100 μM leupeptin, 1 U/mL benzonase) and lysed on ice via sonication. The lysates were clarified by centrifugation and contaminants were precipitated with 5% (V/V) perchloric acid and removed by an additional centrifugation step. The supernatant was then neutralized to pH 7 with neat triethanolamine, dialyzed against equilibration buffer (20 mM HEPES pH 7.4, 0.5 mM PMSF, 50 μM benzamidine, 100 μM leupeptin), and loaded onto a HiTrap SP FF column (Cytiva). CH1 was eluted using a 15–30% gradient of elution buffer (equilibration buffer + 1 M NaCl). Because CH1 lacks aromatic residues, concentration was determined via BCA assay.

Protein labeling with AlexaFluor488-maleimide (AF488, ThermoFisher Scientific, Waltham, MA, USA) or Atto647N-maleimide (ATTO-TEC GmbH, Siegen, Germany) was performed as previously described in Gracia & Polanco et al.^44^ and the degree of labeling was confirmed by absorbance. Following the same protocol, ΔN_t_Tau and K18 were labeled with Tide QuencherTM 2WS-maleimide (TQ2WS, AAT Bioquest, Pleasanton, CA, USA) using the natural cysteine residues at positions 291 and 322.

### Preparation of biomolecular condensates and in vitro phase separation

Heterotypic complex coacervates were prepared at room temperature (22 °C) using a master phase separation buffer. The final assay buffer (PS buffer) contained 10 mM HEPES (pH 7.4), 0.02% (w/V) NaN_3_, 1 mM TCEP (Carbosynth, Compton, UK), 1 mM EDTA (Carbosynth), and 1% (V/V) protease inhibitor cocktail (comprising 100 mM PMSF, 1 mM benzadimide, and 5 μM leupeptin). Unless stated otherwise, the buffer was supplemented with 10% (w/V) polyethylene glycol 8,000 (PEG8k) and 100 mM NaCl. The complete buffer setup was sterilized and clarified through a 0.22 μm syringe filter prior to use.

To eliminate pre-existing oligomeric seeds, individual protein stocks were passed through a 100 kDa molecular weight cut-off (MWCO) centrifugal filter (Amicon Ultra, Millipore, Ireland) immediately before sample preparation. This filtration step was critical to completely remove trace oligomeric species of Tau variants. In the absence of this pre-filtration step, spontaneous amyloid nucleation occurred within the experimental timescale of network consolidation. Upon verification of oligomer removal, amyloid aggregation was not observed under any tested experimental conditions within the 24 h observation window.

Phase separation was initiated in low-binding microcentrifuge tubes by systematically mixing the filtered protein stocks with the master buffer, ensuring that the polyanionic partner (αSyn or N_t_Tau) was invariably added first to control the electrostatic mixing pathway. For crowded conditions utilizing dextran, 10% (w/V) PEG8k was substituted with 15% (w/V) dextran 70,000 (dextran70k). For crowder-free conditions, PEG8k was omitted, and the ionic strength was lowered by adjusting the final NaCl concentration to 20 mM.

To ensure high purity, stock matrices of PEG8k and dextran70k (Biochempeg, Watertown, MA, USA) were dissolved in deionized water and subjected to extensive overnight dialysis against deionized water to eliminate residual salt contaminants in the stock. Following dialysis, the polymer solutions were resuspended, filtered through a 0.22 μm syringe filter, and their final concentrations were precisely quantified via refractometry using a MyBrix Refractometer (Mettler Toledo, Columbus, OH, USA).

### Generation of sonicated αSyn amyloid fibrils

Seed αSyn fibrils were generated by incubating 300 μL of 70 μM αSyn (composed of 69 μM unlabeled protein and 1 μM AF488-labeled αSyn) in PBS (pH 7.4) supplemented with 0.01% (w/V) NaN_3_. The mixture was incubated at 37 °C under continuous orbital shaking at 200 rpm for 7 days. After incubation, the fibril suspension was centrifuged at 9600 ×g for 30 min, and the resulting pellet was resuspended in PBS (pH 7.4). To generate uniform fibrillary seeds, the suspension was sonicated for 1 min (50% duty cycle, 80% amplitude) using a Vibra-Cell VC130 Ultrasonic Processor (Sonics, Newton, USA).

### Charge distribution and NCPR (Net Charge Per Residue) profiles

The net charge per residue (NPCR) diagrams of Tau441 and αSyn were obtained using a custom Python script designed to calculate local charge densities across a sliding window. The protein sequence was discretized into contiguous five-residue segments (“blobs”). The assigned net charge value (*q_i_*) for a given target residue at position *i* was defined as the moving average of the formal charges of the residue itself and its two flanking residues on either side.

### Condensate variable-stringency dissolution assays and quantitative stabilization analysis

To build the phase plots of each system and evaluate the time-dependent network stabilization and aging kinetics of the biomolecular assemblies, phase separated mixtures were incubated at room temperature (22 ± 1 °C) and at designated aging time points, 5 μL aliquots were sampled from the reaction mixture and transferred onto glass-bottom 96-well plates (#1.5H, Ibidi GmbH), covered with adhesive foil, and immediately imaged (within 5 minutes) for the analysis of droplet number and size.

The phase separated samples were prepared (10 μL for each condition for phase plot experiments, 200 μL for kinetic electrostatic challenges) containing specified concentrations of the protein partners (typically 50 μM of Tau variant and 50 μM αSyn). To enable fluorescence visualization, the mixture was doped with 1 μM of a fluorophore-labeled protein (typically AF488-labeled αSyn) in PS buffer. Samples were incubated in 0.2 mL PCR tubes mounted on a benchtop rotating wheel spinning parallel to the longitudinal axis of the tubes; this rotational configuration was maintained continuously to prevent gravity-driven sedimentation of condensates and air bubble formation (which could speed amyloid nucleation in the diluted phase).

For electrostatic challenges, at the different designated time points, condensate stabilities were evaluated via a 1:1 (V/V) variable-stringency chemical dissolution challenge and compare the results with an undiluted 5 μL control sample over time. A 5 μL aliquot harvested from the aging reaction was immediately mixed with 5 μL of a specific challenge buffer to establish two distinct chemical stringencies. Mild challenge (moving upwards in the phase diagram shown in Fig. 1d), where aliquots were mixed with a buffer comprising 10 mM HEPES (pH 7.4), 10% (w/V) PEG8k, 1 M NaCl, and 0.02% (w/V) NaN_3_. This condition disrupts simple non-specific electrostatic networks while preserving the crowding-induced thermodynamic drive. Strong challenge (moving upwards and to the left in the phase diagram shown in Fig. 1d), where aliquots were mixed with the same buffer as in the mild challenge but omitting the PEG crowder to promote further dissolution.

High-resolution three-dimensional (3D) z-stacks (Δz = 5 μm) were immediately acquired on an inverted microscope (Leica DM6000B, Leica Microsystems, Germany) equipped with a spinning-disk confocal scanner (X-Light V2, CrestOptics, Italy) operating with a Nipkow spiral array disk architecture (50/250 μm aperture/spacing). Fluorescence excitation was supplied with a high-power multiline solid-state laser illuminator (89-North LDI-7, 1000 mW) using the 470, 555 and 670 nm laser lines routed through a multiband ZT 405/470/555/640x dichroic filter wheel. Emitted photons were collected using either a 40X dry objective (matrix of 9 independent z-stacks of 10 optical slices each, field of view - FOV: 330×330 μm) or a 63X Fluotar oil-immersion objective (matrix of 18 independent z-stacks of 10 optical slices each, FOV: 211×211 μm). The emission signal was filtered through an ET 525/50 filter (AF488, FITC), 600/50 nm (AmyTracker 630) or ET 700/75 nm filter (Atto647N) bandpass filters. For two-color fluorescence condensate mixing assays of K18:αSyn, a multiband emission filter was used for a simultaneous acquisition of green (AF488) and red (Atto647N) channels. Confocal micrographs were captured on a back-illuminated widefield pco.edge 4.2 bi UV USB sCMOS camera (2048 × 2048 pixels, physical pixel size: 6.5 × 6.5 μm²) using a fixed exposure time of 100 ms to ensure quantitative comparison across all time points. Spatial variations in total imaging area between objective configurations were mathematically normalized during post-processing to ensure accurate cross-sample volumetric comparisons.

Morphometric analysis and droplet segmentation were performed on individual optical slices within ImageJ (v1.52v, NIH, USA) to prevent projection artifacts. A localized binary thresholding mask was applied to each slice, where the dynamic intensity threshold, *I*_Thr_, was defined by local background statistics according to the relation:

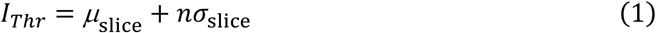

where *μ*_slice_ and *σ*_slice_ represent the mean fluorescence intensity and corresponding standard deviation calculated across that specific optical slice, respectively. A threshold factor of *n* = 4 was used for all phase-plot analyses. For kinetic assays, *n* was adjusted between 4 and 9 on an experiment-by-experiment basis to account for differences in image intensity and signal-to-noise characteristics; however, the selected value was kept constant for all images within the same experiment. In all cases, the binary masks generated by thresholding were visually inspected to confirm accurate droplet detection and segmentation prior to quantitative analysis. Objects displaying a cross-sectional area below 0.2 μm^2^ were systematically excluded from the dataset to eliminate pixel noise and sub-resolution molecular clusters. From these segmented masks, the time-dependent evolution of condensate abundance, area distributions, and dissolution resistance profiles were extracted.

### Determination of the characteristic times of phase-separation and network consolidation

The droplet size distribution was modeled using a log-normal probability density function, defined as:

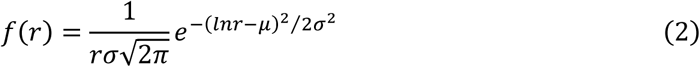

where *r* is the droplet radius, while *μ* and *σ* are the mean and standard deviation of the logarithm of the droplet size, respectively. By fitting the experimental droplet size distributions to this function, *μ* and *σ* were determine at various time points. These parameters were used to track the temporal evolution of the mean droplet size ⟨*r*⟩ = *e^μ^*^+*σ*2⁄2^, the mode (*e^μ^*^+*σ*2^), and the median (*e^μ^*), thereby allowing the extraction of the characteristic timescales.

The temporal evolution of the mean droplet size ⟨*r*⟩ was described using a power-law function, conventional for modeling droplet growth in phase-separation processes:

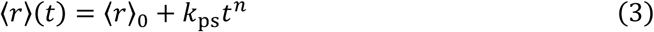

where 〈*r*〉_0_ is the initial mean droplet size, *n* is the growth exponent, and *k*_ps_ is the growth rate constant, such that the characteristic time scale for phase separation, *t*_ps_, is given by *t*_ps_ = k_ps_^−1⁄n^.

To estimate the characteristic timescale of network consolidation or aging, *t*_ag_, we quantified the stabilized fraction of droplets, here referred to as condensate aged fraction (*Φ*_ag_) defined as the ratio of the total condensate area remaining after the dilution challenge *Φ*_ag_ = *A*_after_⁄*A*_before_. The complementary liquid-like fraction of droplets is thus *Φ*_l_ = 1 − *Φ*_ag_. Values of *Φ*_ag_ > 1 arising from measurement noise near full stabilization were set to unity. The temporal evolution of *Φ*_ag_was modeled by a sigmoidal function, as:

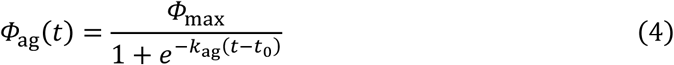

where *Φ*_l_ = 1 − *Φ*_max_ indicates a theoretical maximum value where all droplets are aged, *k*_ag_ is a constant that describes the rate of stabilization, and *t*_0_ is the lag time before the onset of stabilization, such that *t*_ag_ = *t*_0_ + 1⁄*k*_ag_.

### FLIM-FRET data collection and analysis

Nanoscale structural evolution within heterotypic Tau:αSyn coacervates was tracked via time-resolved Förster resonance energy transfer (FRET) using fluorescence lifetime imaging microscopy (FLIM). An Alexa Fluor 488 (AF488) donor was paired with a TideQuencher 2WS (TQ2WS, AAT Bioquest) non-fluorescent dark acceptor to eliminate acceptor-channel fluorescence bleed-through. αSyn was site-specifically labeled with AF488 at position 24 or 122 and incorporated at 5–10 % tracer occupancy. Truncated Tau variants ΔN_t_Tau and K18) were functionalized at native cysteines (Cys291 and Cys322) with TQ2WS-maleimide to constitute 50 % of the total Tau concentration. Samples were loaded into glass-bottom 96-well plates (μ-Plates #1.5H, Ibidi GmbH); K18:αSyn plates were centrifuged at 1,000 x*g* for 5 min immediately post-loading to accelerate early-timepoint droplet sedimentation onto the optical interface without modifying internal condensate morphology.

All FLIM measurements were executed 1 μm above the coverslip surface on an inverted microscope equipped with a 100x Super Apochromat water-immersion objective (NA 1.2, Olympus Life Sciences), a 50 μm confocal pinhole, and a 488/640 nm dichroic mirror (Semrock). Time-correlated single-photon counting (TCSPC) was driven by a pulsed 488 nm laser, with power attenuated between 10 and 100 nW to maintain photon count rates below 500 counts/pixel—preventing detector pile-up anomalies—while ensuring >10^4^ total photons per integrated decay curve. Emitted donor photons were isolated through an ET 520/35 nm bandpass filter, collected via a single-photon avalanche diode detector (SPAD, Micro Photon Devices) at an axial sampling resolution ≥ 0.25 μm/pixel (400 μs pixel dwell time), and compiled using SymphoTime 64 software (PicoQuant GmbH).

For decay analysis, individual condensate regions of interest (ROIs) accumulating >10^4^ total photons were selected, and pixels exceeding 500 counts/pixel were discarded. To bypass instrument response function (IRF) convolutions, a tail-fitting algorithm was initiated at 90% of the peak decay intensity on the trailing edge of the emission peak. The time-resolved fluorescence intensity decay, *I*(t), was fitted to a n-exponential model:

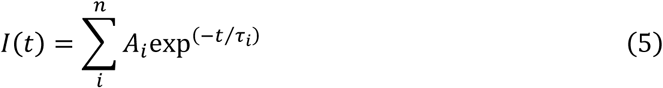

where *A_i_* is the pre-exponential amplitude factor of the *i*-th decay component and *τ_i_* is its discrete lifetime. Optimal fitting (χ^2^ ≈ 1) of FLIM-FRET data required a three-exponential decay model where the fractional contribution *α_i_* of each subpopulation

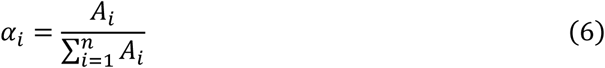

was used to obtain the amplitude-averaged fluorescence lifetime:

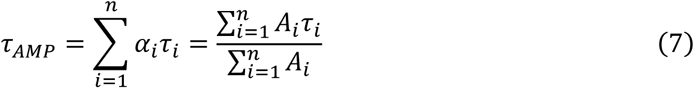

which served as quantitative indicator of FRET efficiency and, thus, donor-quencher proximity.

Typically, 10–20 individual droplet regions of interest (ROIs) were systematically analyzed per experimental condition and longitudinal time point. Replicate measurements were captured across a minimum of two technical replicates per condition, derived from at least three independent biological preparations. For baseline donor-only control experiments (lacking the TQ2WS acceptor), 3–9 droplet ROIs were analyzed per time point across a minimum of three independent biological replicates. Sample sizes were not predetermined using formal statistical power calculations but were selected based on experimental throughput and protein availability while maintaining sufficient numbers to ensure statistical validity. Statistical comparisons were performed using two-tailed Student’s *t-*tests. Homoscedasticity across compared datasets was systematically verified using Levene’s test for equality of variances; equal-variance or unequal-variance (Welch’s)*t-*tests were executed accordingly. The significance of the difference between means is shown as asterisks for the compared pair of data (*p-value < 0.05, **p-value < 0.01, ***p-value < 0.001, ****p-value < 0.0001), and ns means not significant (p-value > 0.05), and the threshold for statistical significance was established at p < 0.05. Quantitative data processing, statistical computing, and plotting were conducted using Microsoft Excel (Office 365), with specific statistical distributions and exact sample sizes (*n*) reported in the respective figure legends.

### Theoretical formulation of the mesoscale series-resistance model

To decouple the multi-scale physicochemical drivers governing biomolecular condensate maturation, we developed a minimal theoretical framework that treats the overall kinetic trajectory as a series-resistance network. The model accounts for three distinct physical mechanisms operating sequentially: interfacial material exchange with the surrounding dilute phase, intra-droplet translational diffusion, and the chemical consolidation of network-forming interactions. Because of the differences between the intrinsic dependences of each process with the condensate size, studying the dependence of the consolidation rates of condensates as a function of their size would allow us to determine the rate limiting step for each system.

We consider a heterotypic system consisting of macromolecular components *A* and *B* that undergo rapid phase separation to establish a dense phase. Within this dense phase, components form transient, reversible intermolecular complexes, denoted as *AB*. Over macroscopic timescales, these under-relaxed native topologies continuously reorganize and consolidate into a stable, hyper-crosslinked network architecture, designated as product *C*. This intra-droplet physical aging process is formalized as:

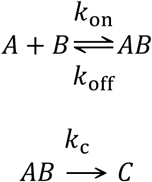

The overall effective rate of network consolidation, *k*_eff_, is determined by the interplay between these transport-and reaction-limited regimes. Operating under the principle of linear addition of characteristic timescales (resistances in series), the total effective maturation time is expressed as:

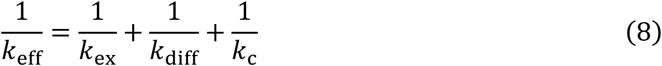

where *k*_ex_ is the rate of interfacial transport, *k*_diff_ is the rate of internal diffusive reorganization, and *k*_c_ is the intrinsic size-independent chemical consolidation rate constant. See Supplementary Material for the comprehensive derivation, radius-dependent scaling formulations, and crossover boundary conditions of each individual resistance component.

## Supporting information

Supplementary Results, Tables and Figures

## SUPPLEMENTARY MATERIALS

Additional results, figures and tables are provided in the Supplementary Information.

## ACKNOWLEDGEMENTS AND FUNDING

This work was supported by the MCIN/AEI/ 10.13039/501100011033 and “ERDF A way of making Europe”, by the European Union (Grant number: PID2022-136997NB-I00) to N.C.; by the Basque Government through the BERC 2022-2025 program and the Spanish State Research Agency through BCAM Severo Ochoa excellence accreditation CEX2021-0011 42-S/MICIN/AEI/10.13039/501100011033 to N.M.; the Diputación General de Aragón Pre-doctoral Research Contract 2021 fellowship to D.P.; and the MCIN/AEI/ 10.13039/501100011033 and “ESF Investing in your future”, by the European Union (PRE2023-UZ-11/PID2022-136997NB-I00) to M.M.

## AUTHOR CONTRIBUTIONS

N.C. conceived the study and designed the experiments. D.P., K.P., A.M and M.M. performed the phase separation assays and dissolution experiments. D.P. and N.C. conducted the in situ single-condensate FLIM–FRET imaging and analyzed the data. N.M. developed the mesoscale series-resistance mathematical model and conducted the numerical analysis on condensates evolution. D.P. and N.C. analyzed the data. D.P., N.M. and N.C. interpreted the results, and framed the non-equilibrium relaxation mechanisms. N.C. wrote the initial draft of the manuscript with input from all authors. N.C. secured funding and supervised the project. All authors reviewed, edited, and approved the final version of the manuscript.

