## Supplementary Results, Tables and Figures for "Rheostatic Network Consolidation Drives Physical Aging in Biomolecular Condensates": Polanco et al_ SI.pdf

### Supplementary Tables

**Supp. Table 1 | Primers used for the generation of new protein variant used in this study.** Other variants were described in Hoyer *et al.*[1] and Gracia & Polanco *et al.*[2]

| Original construct | Target construct | Sense | Primer |
|---|---|---|---|
| Tau441 | $\Delta N_t$Tau | Fw | CTTTAAGAAGGAGATATACATATGATCGCCACACCGCGG |
| | | Rv | CATATGTATATCTCCTTCTTAAAGTTAAAC |
| Tau441 | $N_t$Tau | Fw | GGCTGATGGTAAAACGAAGTGATCAGGCCCCTGGGGCG |
| | | Rv | CTTCGTTTTACCATCAGCCCCCTT |
| Tau441 | AggDef-Tau | Fw | CCGGGAGGCGGGAAGCCAGTTGACCTGAGCAAGGTGACCT |
| | | Rv | CTTCCCGCCTCCCGGCTGG |
| Tau441 | *Δ(1-243)Tau* | Fw | CTTTAAGAAGGAGATATACATATGCAGACAGCCCCCGTGC |
| | | Rv | CATATGTATATCTCCTTCTTAAAGTTAAAC |
| *Δ(1-243)Tau* | K18 | Fw | GCCCCAGGGGCCTGATCATTCAATCTTTTTATTTCCTCCG |
| | | Rv | TGATCAGGCCCCTGGGGCGGTCA |
| αSyn | AggDef-αSyn | Fw | TGACAAATGTTGGAGGAGCAGTGGAGGGAGCAGGG |
| | | Rv | CACTGCTCCTCCAACATTTGT |

### Supplementary Results

#### Validation of macromolecular crowding agents in phase separation and aging kinetics of αSyn electrostatic condensates

Due to the weak, transient nature of the interactions governing between αSyn and Tau complex coacervation, the formation of condensates under diluted conditions in vitro, at low-micromolar concentrations occurs spontaneously under low ionic strength (20 mM NaCl) in the absence of additives (Supp. Fig. 1). Elevating the ionic strength to physiological levels (100–150 mM NaCl) compresses the Debye screening length, screening these long-range electrostatic attractions and shifting the critical coacervation concentration. To systematically interrogate phase separation and subsequent physical aging under physiologically relevant conditions, inert macromolecular crowding agents—specifically polyethylene glycol (PEG8k) and neutral polysaccharides (dextran70k)—were introduced to mimic intracellular excluded volume effects and amplify the thermodynamic activity of the polypeptide chains.

To verify that the progressive physical aging and network consolidation documented herein represent intrinsic relaxation behaviors encoded within the αSyn/Tau sequence architecture rather than microenvironmental artifacts, we systematically evaluated crowding effects across three orthogonal axes.

First, the addition of moderate weight fractions of either PEG8k or dextran70k successfully restored complex coacervation at physiological salinity (Supp. Fig. 1), indicating that phase separation is driven by generic macromolecular volume exclusion rather than specific crowder-protein coordination.

Second, time-resolved fluorescence tracking demonstrated that PEG partitions homogeneously between the continuous dilute phase and the dense coacervate phase, maintaining structural equilibrium at late maturation times (Supp. Fig. 2a). This uniform spatial distribution confirms that crowding operates natively by increasing the effective protein concentration and modulating local water solvation dynamics[3,4], rather than acting through segregative or active depletion-flocculation mechanisms. Furthermore, the partitioning profile of PEG was entirely unaffected by the physical aging of the coacervate network or by exposure to high-stringency chemical dissolution challenges (Suppl. Fig. 2a,b), ruling out time-dependent, depletion-mediated steric compression as the source of droplet stabilization and proving that network maturation is an autogenous property of the protein system.

Third, to explicitly rule out the possibility that structural relaxation and network consolidation are driven by crowder-induced osmotic pressure, we monitored the rate of condensate stabilization across a wide titration matrix ranging from 2.5% to 10% PEG8k (Supp. Fig. 2c). While modulating the crowder weight fraction significantly alters the initial thermodynamic phase boundaries and droplet yield (which were compensated by varying protein concentrations), the internal network stabilization kinetics remained similar across the entire concentration spectrum. This decoupling demonstrates that while macromolecular crowding provides the thermodynamic drive necessary to cross the phase boundary, it does not govern subsequent network consolidation in our experimental systems. Together, these control data validate our reconstituted biophysical platform, confirming that the progressive, sequence-encoded stabilization observed in heterotypic αSyn:Tau condensates is driven entirely by intrinsic, regiospecific electrostatic rearrangements and topological optimization within the internal polymer network rather than by external crowder interference.

**Validation of the regiospecificity of αSyn-Tau interactions during condensate aging**

Independence of network stabilization from fluorophore labeling position

To ensure that the observed network strengthening within the condensed phase was driven by regiospecific sequence-encoded protein-protein interactions rather than fluorophore-induced steric or hydrophobic artifacts, we evaluated the impact of donor positioning. Primary experiments utilized αSyn labeled with Alexa Fluor 488 (AF488) via an engineered cysteine at its negatively charged C-terminus (N122C) — a region known to engage in electrostatic contacts with Tau. We generated an alternative control variant featuring the fluorophore at the distal, amphipathic N-terminus (Q24C), positioning the dye far from the primary Tau-interacting domain (Supp. Fig. 6a).

Using this N-terminally labeled αSyn (Q24C) variant, we repeated the variable-stringency stability challenges at sequential time points during the physical aging of heterotypic PRD311:αSyn coacervates. The progressive development of resistance to both the mild challenge (1:1 dilution in 1 M NaCl, 10% w/V PEG) and the strong challenge (1:1 dilution in 1 M NaCl) was qualitatively and kinetically identical to the profiles obtained using the C-terminally labeled αSyn (N122C) counterpart (Supp. Fig. 6c). These data confirm that fluorophore attachment position does not artificially modulate macroscopic condensate stabilization, network consolidation, or internal aging kinetics.

Regiospecific intermolecular compaction probed by FLIM-FRET

To map the nanoscale structural contributions of distinct αSyn domains to the evolving condensate architecture, we executed time-resolved fluorescence lifetime imaging microscopy (FLIM)–FRET experiments using the AF488-labelled N122C or Q24C αSyn variants as reporters (Supp. Fig. 6a). We hypothesized that if network consolidation is driven by localized electrostatic interactions, structural compaction would manifest predominantly at position 122 (within the negatively charged C-terminus), while position 24 (at the distal N-terminus) would remain spatially segregated from the positively charged Tau stickers (where the TQ2WS acceptor is localized) and, consequently, less sensitive to aging kinetics. Fluorescence lifetime trajectories were recorded immediately following phase separation (0 h) and after extended incubation (4 h for K18:αSyn and 10 h for $\Delta N_t$Tau:αSyn). In these assays, unlabeled wild-type (WT) αSyn was supplemented with 5–10 % of either the 122 or 24 AF488-αSyn reporter variant, while the TQ2WS-Tau variant was introduced at 50 % of the total Tau concentration to maximize the spatial encounter probability between donor and acceptor pairs.

In equimolar $\Delta N_t$Tau:αSyn coacervates (50:50 μM), labeling at position 24 yielded no detectable reduction in donor fluorescence lifetime upon the addition of the TQ2WS-labeled Tau acceptor, tracking identically with quencher-free control condensates (Supp. Fig. 6d). Furthermore, during a 10 h incubation period, the donor lifetime at position 24 remained strictly invariant, whereas position 122 exhibited a pronounced, time-dependent reduction in lifetime. These observations demonstrate that intermolecular network reinforcement and progressive macromolecular compaction in $\Delta N_t$Tau:αSyn condensates are highly region-specific phenomena localized to the negatively charged C-terminal domain of αSyn.

A distinct baseline behavior emerged in equimolar K18:αSyn condensates (50:50 μM). In this framework, probing position 24 revealed a measurable decrease in baseline fluorescence lifetime upon quencher addition, indicating a higher native spatial proximity of the αSyn N-terminus to the K18 chain than to the longer $\Delta N_t$Tau counterpart (Supp. Fig. 6d). This spatial contraction is readily explained by the significantly smaller sequence length and restricted hydrodynamic volume of K18 (129 residues) relative to $\Delta N_t$Tau (292 residues). The truncated K18 backbone shrinks the macroscopic dimensions and internal mesh size of the dense phase, minimizing the steric volume occupied per Tau chain and elevating the spatial density of quencher-bearing sticker motifs. This structural confinement naturally guides the donor-labeled N-terminus of αSyn (residue 24) within the Förster radius of the TQ2WS acceptor even in the absence of a discrete, high-affinity binding locus.

Crucially, despite this enhanced baseline proximity, position 24 exhibited no significant change in fluorescence lifetime between 0 h and 4 h of condensate aging—the exact timescale required to reach maximum network stability (Main Text Fig. 2e), whereas position 122 showed a robust, time-dependent lifetime decay over the same interval (Supp. Fig. 6d). Notably, even in the absence of an acceptor quencher, the baseline fluorescence lifetime at position 24 remained consistently higher than at position 122, revealing that the C-terminus of αSyn natively resides in a significantly more compact and dehydrated local microenvironment within the coacervate.

Taken together, these results support a model wherein condensate stabilization is not uniformly distributed along the primary sequence of αSyn, but emerges through localized, electrostatically driven interactions involving its C-terminal domain. While the K18:αSyn framework features a denser overall network that permits closer non-specific proximity at the N-terminus of αSyn than in $\Delta N_t$Tau:αSyn, the temporal strengthening across both systems remains selectively mediated by the C-terminus of αSyn. These data reinforce the conclusion that regiospecific, charge-complementary contacts dictate the internal structural consolidation and aging kinetics of Tau:αSyn condensates.

### Mathematical derivation and kinetic regimes of the mesoscale series-resistance model

#### Microscopic formulation of individual kinetic resistances

To establish how droplet size scales with network stabilization kinetics, the total effective stabilization rate ($k_{\mathrm{eff}}$) is partitioned intro three physical mechanisms operating sequentially:

First, the exchange of components between the dense droplet and the continuous dilute phase is governed by interfacial transport. Because the interfacial molecular flux scales with the droplet surface area ($A \propto R^2$) while the total protein mass scales with the droplet volume ($V \propto R^3$), the effective volumetric exchange rate, $k_{\mathrm{ex}}$, decreases monotonically as a function of the droplet radius ($R$):

$$k_{\mathrm{ex}} = \frac{3\kappa}{R} \quad (9)$$

where $\kappa$ represents the interfacial permeability coefficient, associated with the local thermodynamic barrier of the phase boundary.

Following interfacial entry or local density fluctuations, polymer chains must diffuse and spatially reorganize within the crowded, viscoelastic environment of the dense phase to achieve productive, complementary alignment. The characteristic timescale for internal diffusive transport scales quadratically with droplet size, yielding a size-dependent diffusion rate constant ($k_{\text{diff}}$) given by:

$$k_{\text{diff}} = \frac{\pi^2 D_{\text{eff}}}{R^2} \tag{10}$$

where $D_{\text{eff}}$ is the effective intra-droplet diffusion coefficient of the macromolecular assembly. Consequently, internal diffusion acts as a profound kinetic bottleneck in large or highly crowded droplets.

Upon successful spatial encounter and local topological alignment, the sticker motifs undergo a chemical consolidation step to permanently strengthen network connectivity. This final molecular relaxation process is strictly size-independent and is governed by an intrinsic activation energy barrier. To interpret $k_{\text{c}}$ in light of non-equilibrium network reconfiguration, we can formalize this step using transition state theory:

$$k_{\text{c}} = \nu_0 \exp\left(-\frac{E_{\text{a}}}{k_{\text{B}}T}\right) \tag{11}$$

where $\nu_0$ is the intrinsic attempt frequency, $E_{\text{a}}$ represents the free energy barrier, analogous to the flow activation energy, $E_{\text{a}}$, that governs intermolecular unbinding and cooperative rebinding within the condensate mesh that has been reported in other biomolecular condensates[5], $k_{\text{B}}$ is the Boltzmann constant, and $T$ is the absolute temperature.

Variations in $k_{\text{c}}$ across distinct protein variants mirror alterations in the fundamental sequence-level sticker grammar rather than geometric or macroscopic transport variations.

<u>Analysis of dominant transport- vs chemical reaction-limited regimes</u>

By plotting the experimental aging rate, defined as $k_{\text{eff}}$, against the droplet radius ($R$), the series-resistance model identifies distinct, size-dependent kinetic regimes dictated by the relative magnitudes of $\kappa$, $D_{\text{eff}}$, and $k_{\text{c}}$:

**Transport-Limited Regime (Fast $k_{\text{c}}$):** When intrinsic molecular consolidation rate is rapid, $k_{\text{eff}}$ is entirely controlled by material transport limitations. For small droplets below a critical threshold ($R << R_x$), the system is surface-exchange-limited, exhibiting an $k_{\text{eff}} \propto R^{-1}$ scaling. Conversely, for large droplets ($R >> R_{\text{x}}$), internal translational diffusion becomes rate-limiting step ($k_{\text{eff}} \propto R^{-2}$). The transition between these regimes occurs at a characteristic crossover radius ($R_x$), defined by setting $k_{\text{ex}}$ = $k_{\text{diff}}$

$$R_{\text{x}} = \frac{\pi^2 D_{\text{eff}}}{3\kappa} \tag{12}$$

**Reaction-Limited Regime (Slow $k_{\text{c}}$)**: If the energetic barrier $E_{\text{a}}$ is high, chemical consolidation is slow compared to both interfacial exchange and diffusion. In this regime, $k_{\text{eff}}$ remains size-independent ($k_{\mathrm{eff}} \approx k_{\mathrm{c}}$) in smaller droplets. However, due to the power-law dependence of transport resistances, intra-droplet diffusion inevitably overtakes the reaction rate as the rate-limiting bottleneck once a droplet grows past a critical geometric threshold.

Measuring $k_{\mathrm{eff}}$ as a function of droplet radius successfully decouples the intrinsic interaction energetics from macroscopic transport contributions. While the diffusion-limited regime reflects long-range macromolecular motion sensitive to local crowding and droplet size, the reaction-limited regime maps directly to the sequence-encoded unbinding/rebinding landscape of the network.

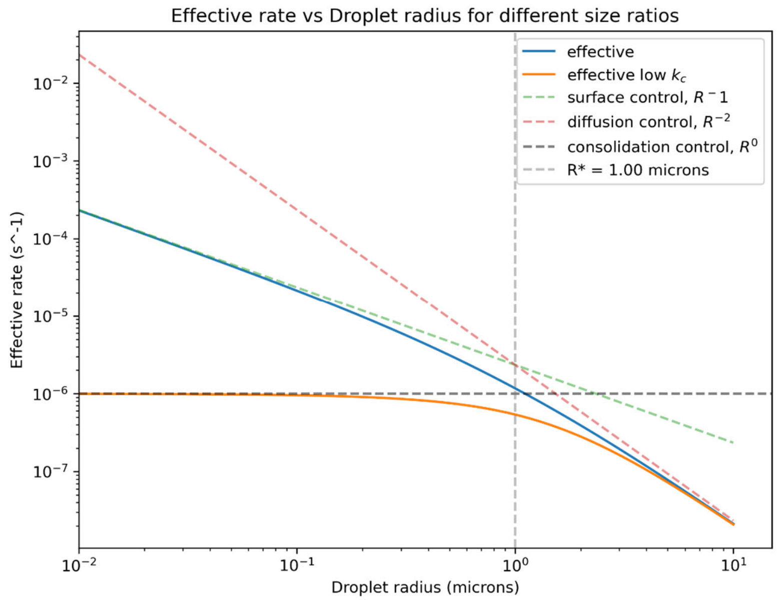

**Size-dependent stabilization kinetic of biomolecular condensates via the series-resistance model**. The effective rate of network stabilization ($k_{\mathrm{eff}}$) is determined by the dynamic interplay between interfacial material exchange ($k_{\mathrm{ex}}$), internal diffusion within the droplet dense phase ($k_{\mathrm{diff}}$), and intermolecular network consolidation ($k_{\mathrm{c}}$). In regimes where chemical consolidation ($k_{\mathrm{c}}$) is fast, $k_{\mathrm{eff}}$ is governed entirely by macromolecular transport processes: interfacial exchange with the continuous dilute phase limits the rate in small droplets ($k_{\mathrm{eff}} \propto R^{-1}$), whereas internal diffusion limits the rate in larger droplets ($k_{\mathrm{eff}} \propto R^{-2}$). The transition between these surface-limited and diffusion-limited transport regimes occurs at a characteristic crossover radius ($R_{\mathrm{x}}$). In regimes where $k_{\mathrm{c}}$ is slow, the intrinsic interaction consolidation process dominates and yields size-independent kinetics in small droplets. However, with increasing droplet size, internal translation diffusion inevitably becomes the rate-limiting bottleneck due to its quadratic dependence on the droplet radius ($R^{-2}$).

Mapping the scaling framework in systems with well-dispersed vs continuous stickers

This scaling framework provides a quantitative basis for identifying dominant kinetic regimes across diverse experimental protein systems. By evaluating the dependence of $t_{\mathrm{ag}}$ for different droplet radius ($R$), we mapped the aging dynamics of $\Delta N_t$Tau:αSyn (50:50, 20:20, and 10:10 μM), and $\Delta N_t$Tau:$N_t$Tau (50:50 μM) (Figures 5 in the Main Text and Supp. Fig. 8).

To obtain the rates of aging as a function of droplet sizes, we used the histograms of condensates obtained in the variable-stringency dissolution assays, where the condensates at different maturation times were subjected to two independent dissolution conditions and the fraction of condensates remaining was determined. The mild and strong challenge conditions probe distinct aging states along a continuous liquid-to-gel consolidation landscape (Figure 6a-b in the Main Text) rather than discrete material phases. Mild challenges predominantly interrogate shallower, weakly consolidated network states and exhibit relatively high $k_{\mathrm{eff}}$ values (Supp. Fig. 8a). Strong challenges, by contrast, probe more deeply consolidated regions of the energy landscape, where increased network connectivity and reduced relaxation rates result in lower $k_{\mathrm{eff}}$ (Supp. Fig. 8b).

More generally, the two conditions shown here should be viewed as representative snapshots of a continuum of network consolidation states that can be accessed by varying challenge strength, aging time, composition, or environmental conditions. Across this continuum, the same series of kinetic resistances—molecular exchange with the surrounding phase, internal diffusive rearrangements, and network consolidation—are expected to govern the observed aging dynamics. The relative contribution of each resistance, however, shifts as the system descends deeper into the gel free-energy basin (Fig. 6a-b).

Correspondingly, the magnitude of $k_{\mathrm{eff}}$ decreases and the sensitivity to droplet size increases as condensates become progressively more consolidated. The concentration dependence observed for the $\Delta N_t$Tau:αSyn systems is consistent with this picture: higher concentrations generally exhibit stronger size dependence, indicative of a larger contribution from transport- and relaxation-limited processes, whereas lower concentrations display weaker radius dependence and larger effective rates, consistent with shallower consolidation states and more efficient network reorganization. Together, these results support a unified framework in which condensate aging is governed by the same underlying resistance network across a spectrum of consolidated states, with the depth of consolidation determining both the absolute timescale of aging and the dominant kinetic bottlenecks.

### Supplementary Figures

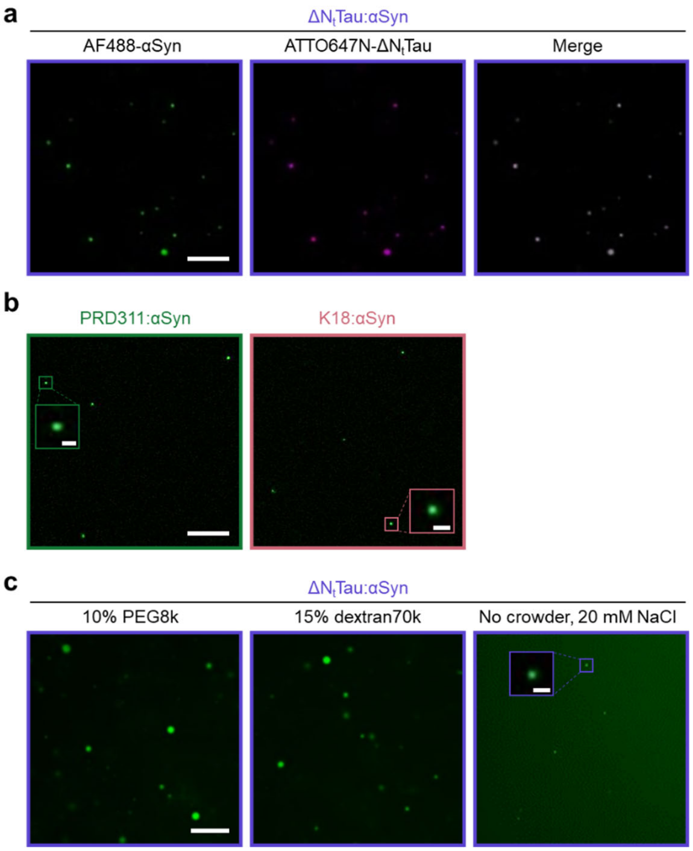

**Supp. Fig. 1 | Complex coacervation of αSyn with the three Tau constructs. a**, Confocal (CF) microscopy images of $\Delta N_t$Tau:αSyn condensates at 50:50 µM containing 1 µM Atto647N-$\Delta N_t$Tau (magenta) and 1 µM AF488-αSyn (green), formed in PS buffer and imaged after 1 h incubation. **b**, CF microscopy images of PRD311:αSyn (left) or K18:αSyn (right) condensates at 50:50 µM containing 1 µM AF488-αSyn, formed in PS buffer and imaged after 1 h incubation. **c**, CF microscopy images of $\Delta N_t$Tau:αSyn condensates at 50:50 µM containing 1 µM AF488-αSyn, formed and incubated for 4 h under varying solution conditions: standard PS buffer (left), PS buffer prepared with 15% dextran70k instead of 10% PEG8k (center), and PS buffer in the absence of macromolecular crowder with 20 mM NaCl instead of 100 mM NaCl (right). Scale bars represent 10 µm and apply to all images within the respective panels.

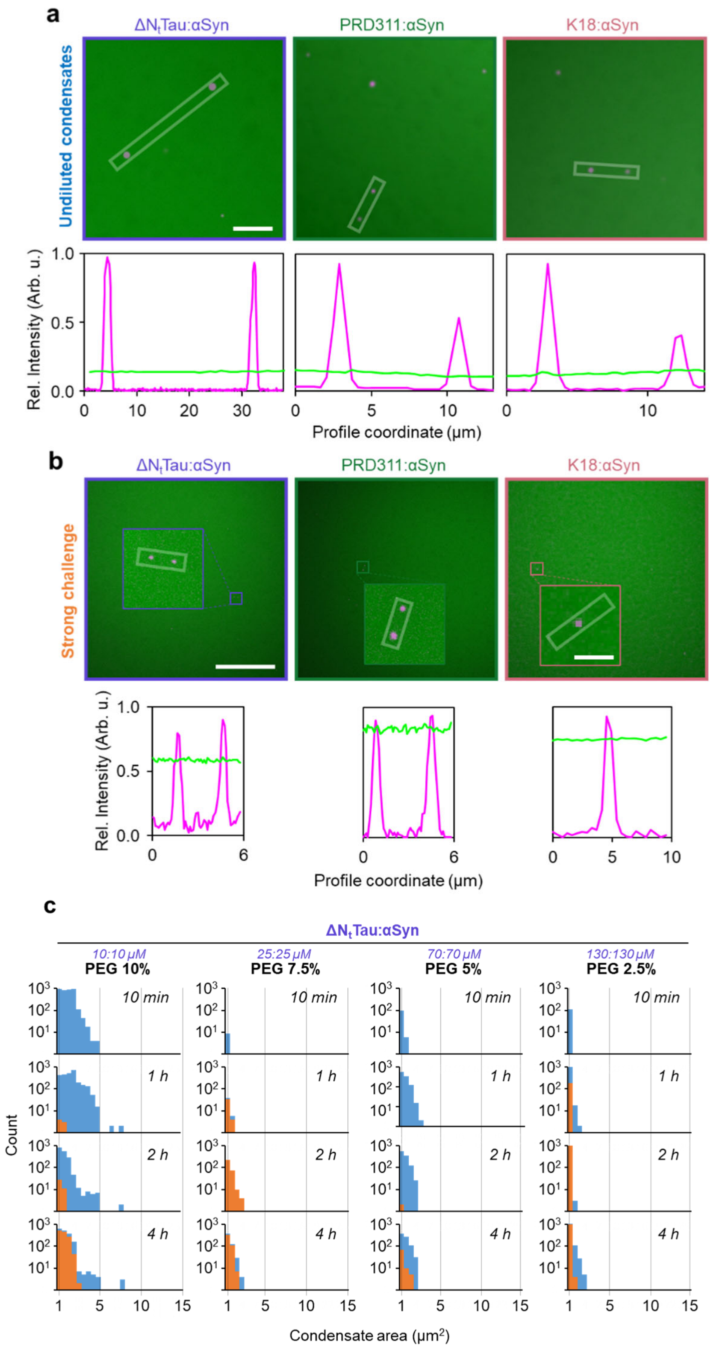

**Supp. Fig. 2 | Macromolecular crowder partitioning and concentration-dependent condensate stability during aging. a-b**, CF microscopy images tracking the aging-dependent partitioning of FITC-PEG8K (green) into $\Delta N_t$Tau:αSyn (left), PRD311:αSyn (middle), and K18:αSyn (right) condensates at 50:50 µM stoichiometric ratio, supplemented with 1 µM Atto647N-αSyn (magenta) as a dense-phase marker. Imaged were acquired after 4 h incubation either before (**a**), or after (**b**) a 1:1 dilution with the high-stringency electrostatic challenge (strong challenge, see Methods). Lower panels display corresponding normalized linear fluorescence intensity profiles for FITC-PEG8K (green) and Atto647N-αSyn (magenta) measured across the representative spatial vectors indicated by dashed white rectangles in the associated CF images, demonstrating homogeneous interfacial partitioning of the PEG crowder within both the continuous dilute and dense coacervate phases across progressive aging states and following high-stringency chemical disruption applied to evaluate network stability. Scale bars represent 10 µm in panel **a**, 50 µm in panel **b**, and 5 µm for the corresponding high-magnification inset zoomed condensates in panel **b**; scale bars apply to all orthogonal images within their respective panels. **c**, Time-dependent network stabilization of $\Delta N_t$Tau:αSyn condensates formed and incubated at varying PEG8K concentrations in PS buffer. Representative droplet size distributions at designated incubation times before (blue histograms) and after (orange histograms) a 1:1 dilution with the strong dissolution buffer are shown for conditions ranging from 2.5 to 10% w/V PEG8k. To maintain relatively homogeneous droplet number density and comparable population statistics across identical microscopy acquisition settings, the total bulk protein concentration was systematically scaled inverse-proportionally to the target volume exclusion (as indicated) to compensate for phase-boundary shifts while maintaining a strict 1:1 heterotypic stoichiometry. The population profiles reveal robust, time-dependent condensate stabilization over a 4 h incubation period with qualitatively invariant aging kinetics across all tested PEG8k regimes. These results indicate that macromolecular crowding does not significantly alter the underlying kinetic network consolidation timescales, confirming that the physical aging of the αSyn–Tau electrostatic network is an intrinsic molecular process driven by the associative polymer chemistry inherent to the protein condensates.

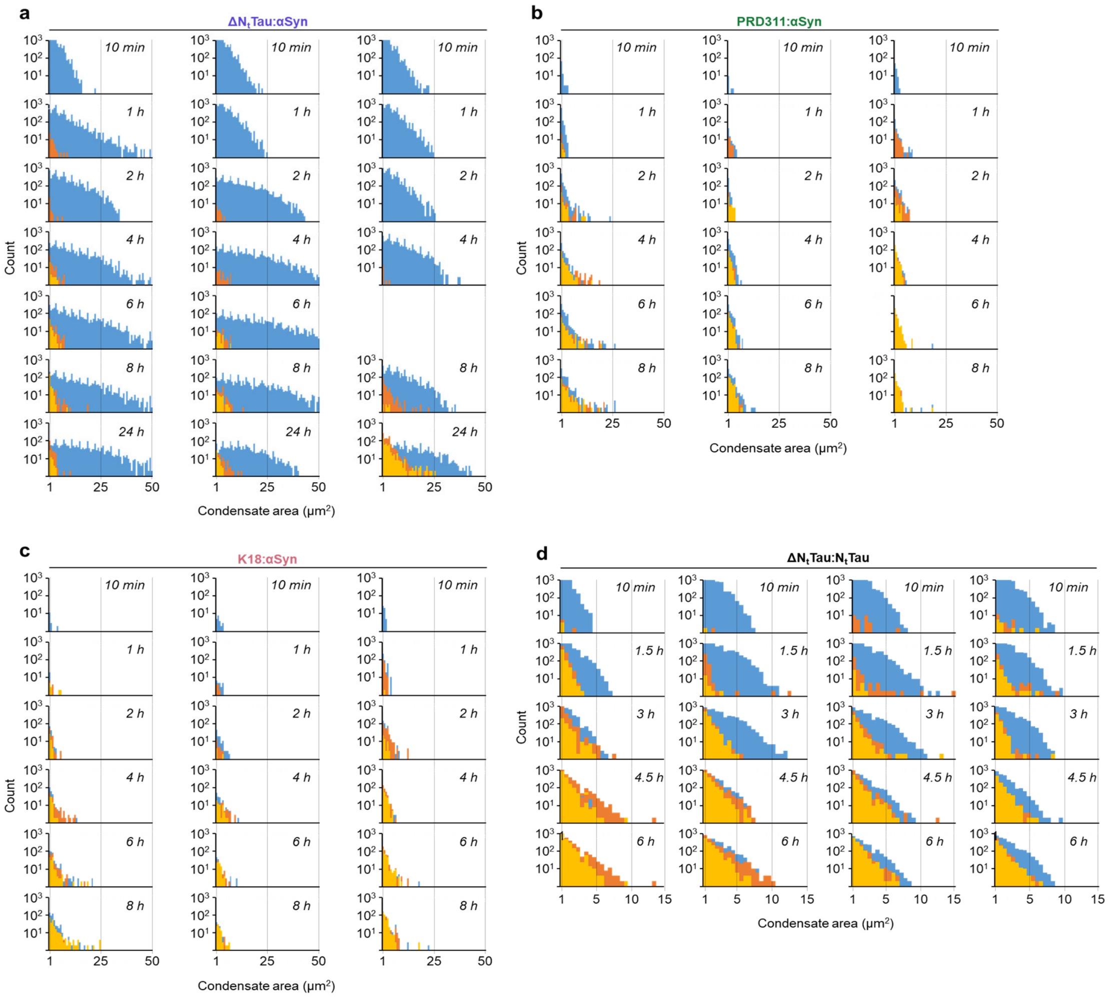

**Supp. Fig. 3 | Replicate droplet size distributions and stability profiles across the main four condensate systems.** Droplet size distributions as a function of aging time for **a**, ΔN$_t$Tau:αSyn, **b**, PRD311:αSyn, **c**, K18:αSyn and **d**, ΔN$_t$Tau:N$_t$Tau condensates formed at 50:50 μM in PS buffer. Population histograms represent unperturbed sample size distributions in standard PS buffer (blue), samples subjected to a 1:1 dilution with the low-stringency challenge buffer (mild challenge, orange), and samples subjected to a 1:1 dilution with the high-stringency electrostatic challenge buffer (strong challenge, yellow) (see Methods for detailed buffer compositions). For each biomolecular system, a single representative experimental replicate is displayed, with each individual longitudinal time point comprising data aggregated from 9 distinct fields of view spanning 10 optical slices per confocal Z-stack.

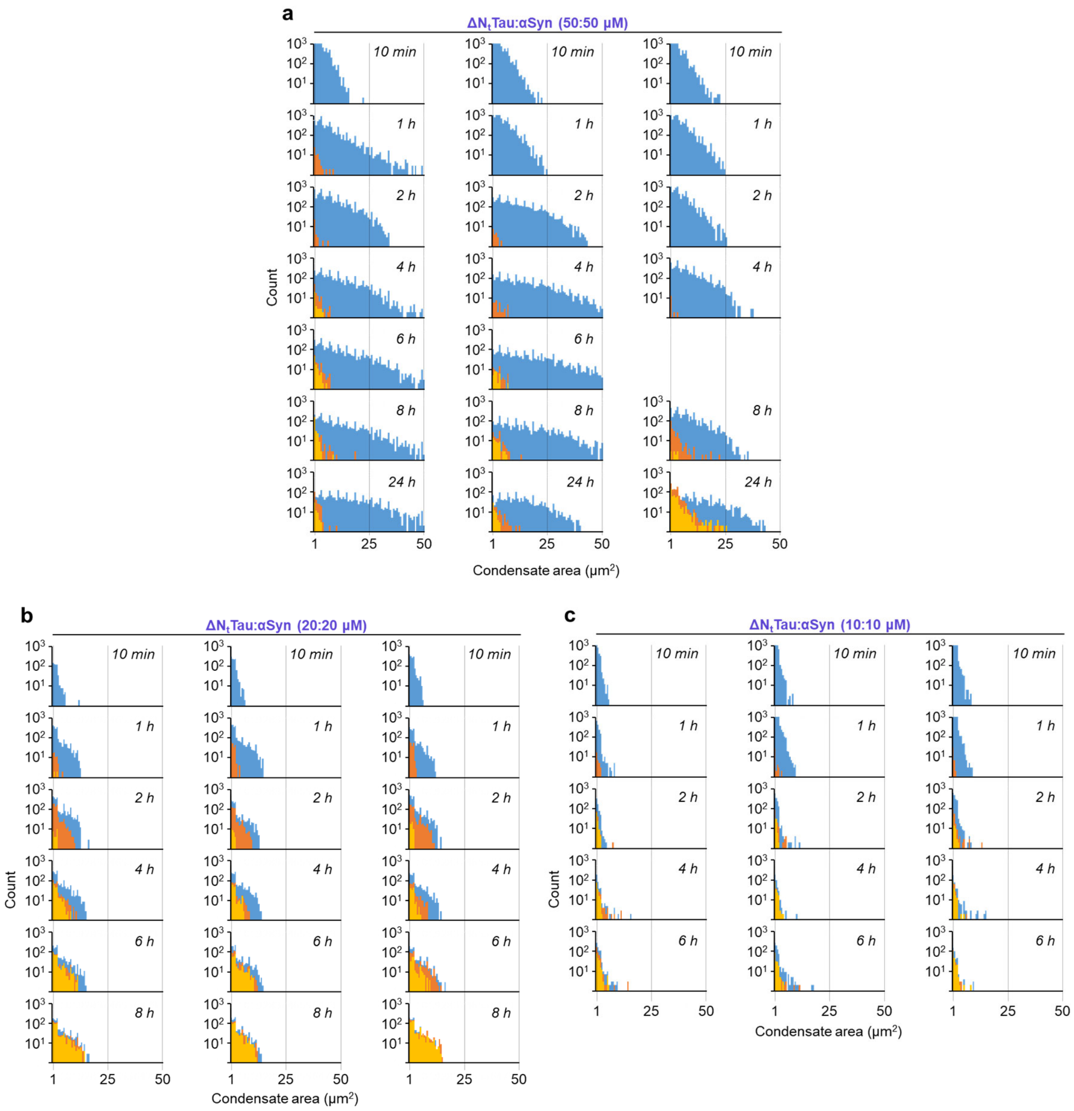

**Supp. Fig. 4 | Replicate droplet size distributions and stability profiles across the three $\Delta N_t$Tau:αSyn concentrations.** Droplet size distributions as a function of aging time for $\Delta N_t$Tau:αSyn condensates formed at **a**, 50:50 µM, **b**, 20:20 µM and **c**, 10:10 µM in PS buffer. Population histograms represent unperturbed sample size distributions in standard PS buffer (blue), samples subjected to a 1:1 dilution with the low-stringency challenge buffer (mild challenge, orange), and samples subjected to a 1:1 dilution with the high-stringency electrostatic challenge buffer (strong challenge, yellow) (see Methods for detailed buffer compositions). For each biomolecular system, a single representative experimental replicate is displayed, with each individual longitudinal time point comprising data aggregated from 9 distinct fields of view spanning 10 optical slices per confocal Z-stack.

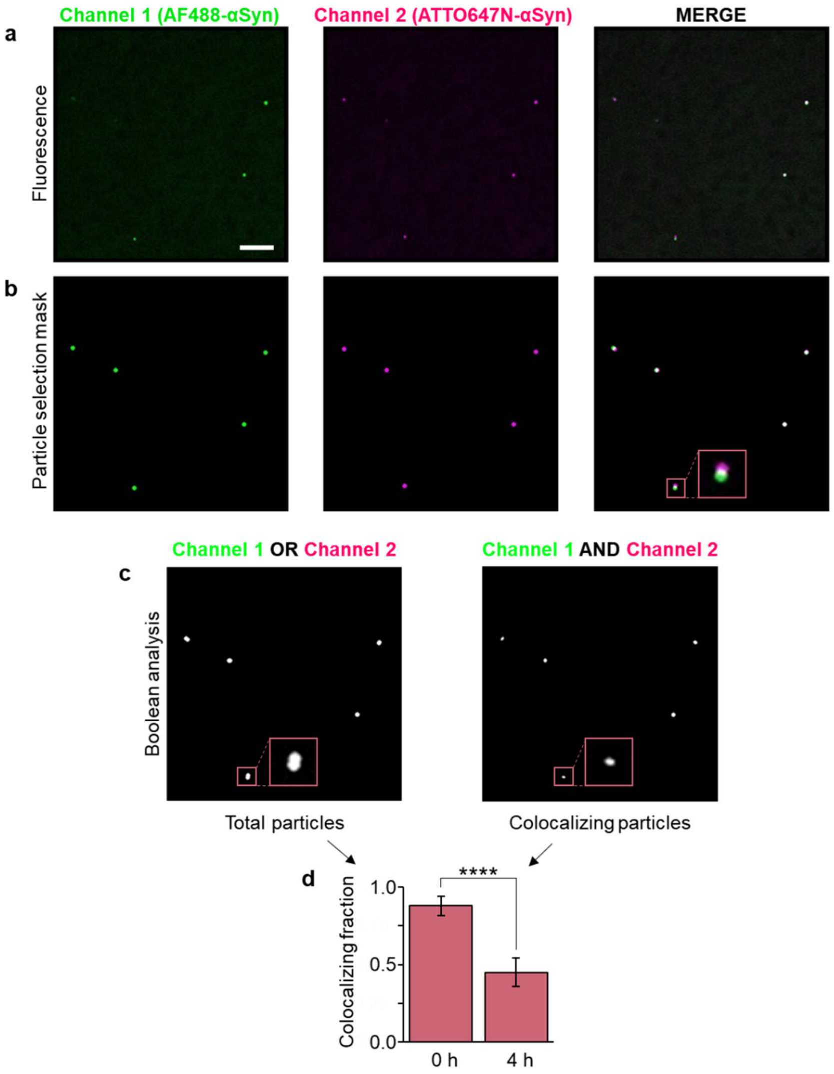

**Supp. Fig. 5 | Image-processing workflow for Figure 3f. a**, A single optical slice of K18:αSyn condensates formed separately at 50:50 µM (fluorescently labelled with either 1 µM AF488-αSyn or 1 µM Atto647N-αSyn) acquired simultaneously by CF microscopy across distinct channels (channel 1: green; channel 2: magenta, Merge: composite) and imaged after 1 hour post-mixing. **b**, The thresholding workflow detailed in the Methods section generates a binary particle selection mask for each individual channel. A zoomed particle containing overlapping signal from both fluorophores is highlighted in pink. Note that particles are not approximated as single mathematical points; instead, their cross-sectional boundaries are preserved and radii estimated to match the physical dimensions of the original condensates. **c**, Boolean-based logical segmentation to quantify conjunction (total, "OR", left image) and intersection events (colocalization "AND", right image). The scalebar corresponds to 10 µm for all images. **d**, Quantification of the colocalizing fraction. For each longitudinal time point, bars represent the mean colocalizing fraction computed from 6 independent fields of view across 10 optical sections each (yielding a total population size of $n$= 526 condensates for 0 h and $n$= 192 for 4 h; error bars denote standard deviations. Statistical significance from a two-tailed Student's $t$-test is indicated by asterisks (*****p-value < 0.05, ******p-value < 0.01, *******p-value < 0.001, ********p-value < 0.0001), where "ns" denotes a non-significant variance (p-value ≥ 0.05).

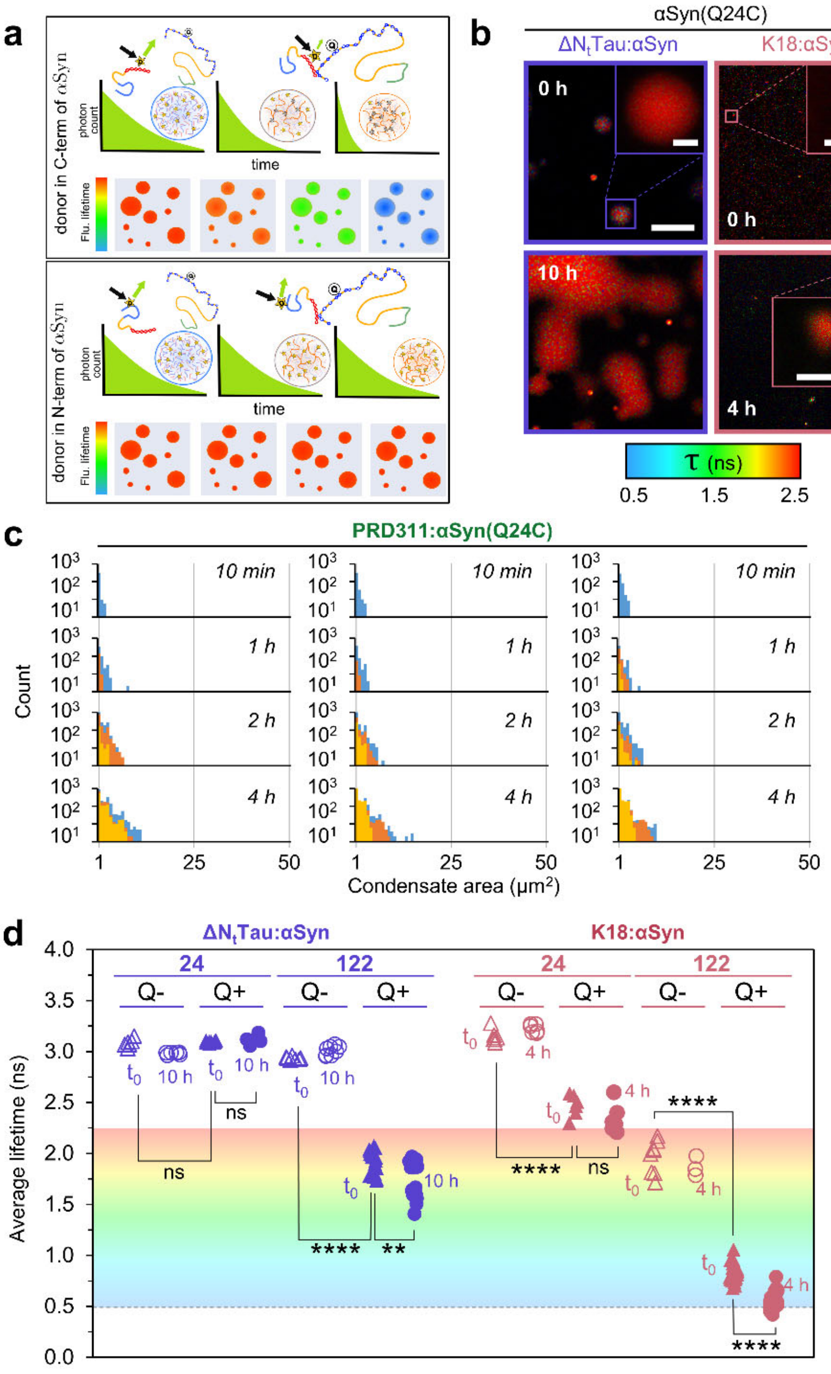

**Supp. Fig. 6 | Effect of αSyn labelling position on aging and FLIM-FRET mapping of region-specific network consolidation. a**, Schematic representation of expected fluorescence lifetime changes for distinct donor-labeling positions in FLIM–FRET assays. In loosely interacting condensates or condensates without acceptor molecule, AF488 remains essentially unquenched, exhibiting an invariant long fluorescence lifetime. Upon physical aging of the coacervates, C-terminally labeled αSyn (upper panel) experiences localized proximity via progressive strengthening of electrostatic interactions with acceptor-bearing TQ2WS-Tau stickers, yielding a systematic, time-dependent reduction in donor lifetime.

Conversely, for N-terminally labeled αSyn (lower panel), the donor and TQ2WS acceptor remain separated beyond the Förster radius, precluding efficient energy transfer and verifying the regiospecificity of the network interactions (see Supp. Results). **b**, Representative FLIM images displaying the intensity-weighted average fluorescence lifetime of AF488-αSyn(Q24C). Purple panels show TQ2WS-ΔN$_t$Tau:αSyn condensates at 0 h (top) and 10 h (bottom). Pink panels show TQ2WS-K18 αSyn condensates at 0 h (top) and 4 h (bottom). The pixel-level pseudocolor scale is bounded between 0.5 and 2.5 ns to match the dynamic range in Figure 4. **c**, Droplet size distributions as a function of aging time for 50:50 µM PRD311:αSyn condensates containing 1 µM N-terminally labeled AF488-αSyn(Q24C). Histograms show unperturbed samples in standard phase-separation (PS) buffer (blue), and samples post-1:1 dilution with either mild (orange) or strong (yellow) challenge buffers (see Methods). Data represent a single experimental replicate aggregated from 9 fields of view across 10 optical planes. **d**, Absolute fluorescence lifetimes of AF488 in FRET-active (Q+) versus donor-only (Q-) control condensates, contrasting AF488-αSyn(Q24C) against AF488-αSyn(N122C). Profiles are plotted for ΔN$_t$Tau:αSyn (purple; 0 h triangles, 10 h circles) and K18 αSyn (pink; 0 h triangles, 4 h circles), using filled (Q+) and hollow (Q-) symbols. Background gradients match the pseudocolor lifetime scale for panels b and d. Statistical significance from two-tailed Student's *t*-tests is indicated by asterisks (*p-value < 0.05, **p-value < 0.01, ***p-value < 0.001, ****p-value < 0.0001), where “ns” denotes p-value ≥ 0.05.

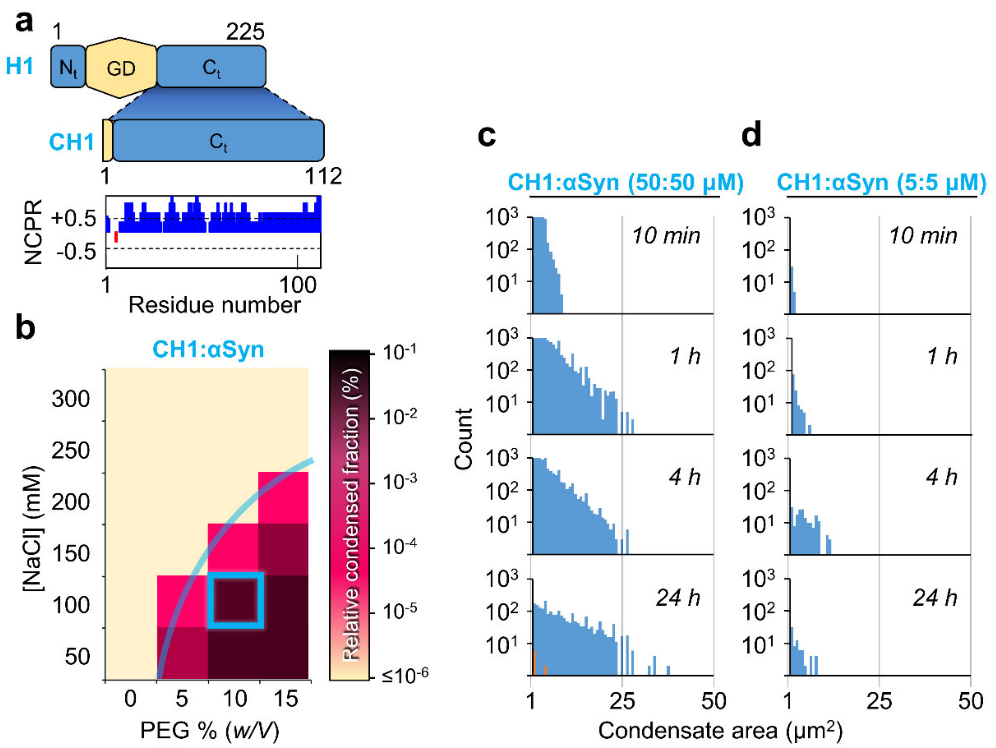

**Supp. Fig. 7 | Complex coacervation of αSyn with the C-terminal domain of histone H1. a**, Schematic of C-terminal domain truncate used in this study (CH1) of chicken histone H1 protein, showing its regions: a remaining of the structured globular domain (residues 1-12, in yellow), and the intrinsically disordered C-terminal domain ($C_t$, in blue). The net charge per residue (NCPR) of CH1 is shown below. **b**, Phase plot showing the relative condensed fraction of CH1:αSyn at 50:50 µM (+1 µM AF488-αSyn) at varying NaCl and PEG8k concentrations (9 fields of 10 slices for each single condition, N=3). Condensates were imaged by confocal microscopy 1 h after phase separation. Color scale indicates the condensed fraction relative to the maximum condensed fraction amongst all the conditions tested. The chosen buffer condition for most of the following experiments (named PS buffer) is highlighted in blue. **c-d**, Size distribution of CH1:αSyn condensates formed at **c**, 50:50 µM or **d**, 5:5 µM, and imaged at each given time point during incubation. Original sample size distributions in PS buffer are shown in blue, and challenged sample distributions are shown in orange (mild challenge) and yellow (strong challenge). A single replica for each protein concentration is shown.

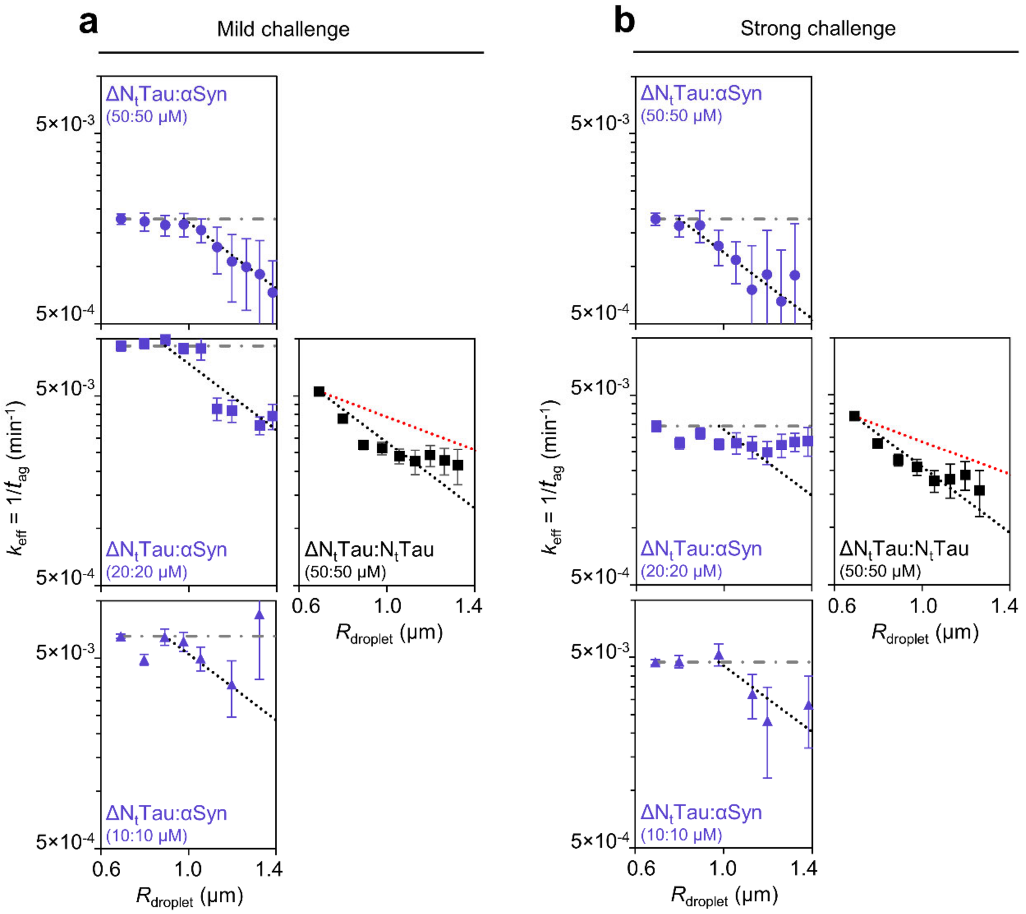

**Supp. Fig. 8 | Effective stabilization rates scale with droplet size under mild and strong challenge conditions. a-b**, Size-resolved effective stabilization rate constants ($k_{\mathrm{eff}}$) plotted as a function of droplet radius ($R_{\mathrm{droplet}}$) for distinct protein systems and stoichiometries under **a**, mild challenge and **b**, strong challenge conditions. Purple symbols correspond to heterotypic ΔN$_t$Tau:αSyn condensates at the indicated concentrations (50:50, circles; 20:20, squares; 10:10 μM, triangles), whereas black symbols correspond to ΔN$_t$Tau:N$_t$Tau condensates at 50:50 μM. Data points represent experimentally determined effective gelation rates obtained from droplet-ageing measurements within individual radius bins. Dotted lines are included as guides to the eye and indicate characteristic scaling behaviors expected for distinct rate-limiting mechanisms: $k_{\mathrm{eff}} \propto R^{-1}$ (red dotted line), consistent with surface-limited processes; $k_{\mathrm{eff}} \propto R^{-2}$ (black dotted line), consistent with diffusion-limited transport; and $k_{\mathrm{eff}} \propto R^{0}$ (gray dashed-dotted line), corresponding to size-independent bulk kinetics. Error bars represent the relative uncertainty in $k_{\mathrm{eff}}$, estimated assuming Poisson-like counting statistics according to $\sigma_{k_{\mathrm{ag}}}/k_{\mathrm{ag}} = 1/\sqrt{N_{\mathrm{drop}}}$, where $N_{\mathrm{drop}}$ is the number of droplets analyzed within each radius bin.
